# *RTL4,* a retrovirus-derived gene implicated in autism spectrum disorder, is a microglial gene that responds to noradrenaline in the postnatal brain

**DOI:** 10.1101/2024.10.23.619930

**Authors:** Fumitoshi Ishino, Johbu Itoh, Ayumi Matsuzawa, Masahito Irie, Toru Suzuki, Yuichi Hiraoka, Masanobu Yoshikawa, Tomoko Kaneko-Ishino

**Author notes:** Research Infrastructure Management Organization, Institute of Science Tokyo, Tokyo113-8510, Japan (Fumitoshi Ishino). Logomix, Inc. Tokyo 104-0053, Japan. Nonprofit Organization Gene Information Bank, Fukuoka 810-0041, Japan (Masahito Irie). Correspondence to: Fumitoshi Ishino and Tomko Kaneko-Ishino.

## Abstract

*RTL4*, a gene acquired from a retrovirus, is a causative gene in autism spectrum disorder. Its KO mice exhibit increased impulsivity, impaired short-term spatial memory, failure to adapt to novel environments, and delayed noradrenaline (NA) recovery in the frontal cortex. However, due to its very low expression in the brain, it remains unknown which brain cells express RTL4 and its dynamics in relation to NA. Here, using knock-in mice carrying endogenous *Rtl4* fused to *Venus*, we demonstrated that *RTL4* is a microglial gene with several important properties. The RTL4-Venus fusion protein was detected as a secreted protein in the midbrain, hypothalamus, hippocampus and amygdala in the postnatal brain. Its signal intensity was high during critical periods of neonatal adaptation to novel environments and was upregulated by various stimuli, such as handling the mice, mild environmental changes and administration of isoproterenol, an agonist of adrenergic beta receptors. It was decreased by anesthesia but was maintained by the administration of milnacipran, an NA reuptake inhibitor. These results suggest that RTL4 intensity depends on the arousal state via NA and is somehow related to the NA reuptake process. *In vitro* mixed glial culture experiments demonstrated that *Rtl4* is a microglial gene and suggested that RTL4 secretion responds rapidly to isoproterenol. Taken together with previous results from *Rtl4* KO mice, microglial RTL4 plays an important role in the NA response and it is likely involved in the development of the NAergic neuronal network in the brain.

## Introduction

Retrotransposon Gag-like (RTL)/sushi-ichi retrotransposon homolog (SIRH) genes are eutherian-specific, with the exception of therian-specific *PEG10*. They were acquired by three independent domestication events of a certain retrovirus-like retrotransposon (*Metaviridae*) during mammalian evolution [1–11]. It is reasonable to assume that they were originally derived from the GAG and POL of a certain extinct retrovirus with a high degree of homology to the sushi-ichi retrotransposon, since the gypsy retrotransposon, which includes the sushi-ichi retrotransposon, is an infectious retrovirus in *Drosophila melanogaster* [12, 13]. Of these eleven genes, at least ten RTL/SIRH genes are known to have specific essential and/or important functions in current eutherian developmental systems, such as the placenta and brain [8, 10]. Among them, *RTL4* (aka *SIRH11* or zinc finger CCHC domain-containing protein 16 (*ZCCHC16*)) has been implicated through extensive screening of patients as a causative gene in autism spectrum disorder (ASD) [14]. Lim *et al.* identified a family with a rare nonsense mutation in X-linked gene *RTL4* (*ZCCHC16*) leading to ASD in a male proband and his male sibling [14]. Consistent with this, we previously demonstrated that *Rtl4* (*Sirh11/Zcchc16*) KO mice exhibit increased impulsivity, decreased attention, poor working memory and reduced adaptation to novel environments [15]. They displayed low noradrenaline (NA) recovery in the frontal cortex, suggesting that the behavioral defects of the *Rtl4* KO mice are somehow related to a dysregulation of the NA system in the brain [15]. Despite its important role in the brain, several critical questions, such as which brain cells express RTL4 protein and when, where, and how it functions in relation to NA, have long remained unelucidated, not least because *RTL4* expression is quite low throughout the body, including the brain [16, 17].

NA-expressing neurons have been reported to play important roles in attention, vigilance, behavioral flexibility and modulation of cognition [18–21], and their activation occurs in concert with the cognitive shifts that facilitate dynamic reorganization of target neural networks, allowing rapid behavioral adaptation to the changing environmental demands [20]. NA is also the most well-documented neurotransmitter in stress experiments. NA has been reported to increase in the brain in response to various types of stress [22], and administration of the β-adrenergic receptor (AR) agonist isoproterenol significantly increases interleukin-1β in the brain [23, 24] and cultured microglia [22, 25]. Most of the secreted NA is reabsorbed by a noradrenaline transporter (NAT, aka norepinephrine transporter, NET) located along the plasma membrane of the presynaptic neuron after it has completed its function of transmitting a neural impulse [26]. Therefore, for adequate maintenance of intracellular NA stores in NA neurons, NAT-mediated reuptake of NA is critical [26]. Milnacipran is a serotonin (5-hydroxytryptamine, 5-HT) -NA reuptake inhibitor (SNRI) that is thought to increase the levels of synaptic 5-HT and NA by inhibiting their reuptake into the neuronal cells, thereby exhibiting antidepressant and anxiolytic effects [27].

Neurotransmitters also play an important role in nervous system development, including the shaping and wiring of the nervous system during critical periods of development [28, 29]. In particular, NA is postulated to be an important regulator of brain development, regulating the development of both NAergic neurons and their target areas during early stages of development, affecting such behaviors as infant attachment leaning, aversion and fear learning, as well as synaptogenesis. Disruptions in this process can alter the trajectory of brain development, leading to long-term and even permanent changes in brain function and behavior later in life [29].

NA signals through three types of ARs, α1, α2 and β, each of which in turn has three subtypes, α1A, α1B, α1D; α2A, α2B, α2C; β-1, β-2, β-3, respectively. All are G protein-coupled receptors, with α1-ARs coupled to G_q_, α2-ARs coupled to G_i/o_ and β-ARs coupled to G_s,_ and have distinct pharmacological properties, molecular structures and signaling pathways [30]. These ARs are present on multiple cell types throughout the CNS, including astrocytes and microglia, as well as neurons [31, 32]. The α1- and β-ARs are present at the postsynaptic site and generally mediate excitatory effects, whereas α2-ARs are present at both the pre- and postsynaptic sites and reduce NA release by decreasing neuronal excitability [33].

Microglia are the primary innate immune cells of the brain and play a central role in the immune response to various pathogens via a variety of Toll-like receptors (TLRs) [34, 35] and/or in the clearance of various pathogen-associated molecular patterns (PAMPs) such as single-stranded (ss)RNA/double-stranded (ds)DNA (viruses), lipopolysaccharide (LPS) (Gram-negative bacteria) and zymosan (fungi) by means of RTL/SIRH proteins [36, 37]. This is accomplished by the rapid capture and degradation-associated activities of RTL5 (aka SIRH8), RTL6 (aka SIRH3) and RTL9 (aka SIRH10), respectively [36, 37]. Moreover, in the neonatal brain microglia are involved in shaping neuronal circuits during development by regulating neurogenesis. They induce filopodia formation by direct contact with neurons and phagocytose supernumerary or unneeded synapses [34, 38, 39]. Therefore, they have been implicated as an important cause of several neurodevelopmental and neuropsychiatric disorders, including ASD [40–43].

In this work, using knock-in (KI) mice carrying a gene that fuses endogenous *Rtl4* with *Venus* (*Rtl4*CV mice) to produce the RTL4-Venus fusion protein at the C-terminus (RTL4CV), we demonstrated that *Rtl4* is a fourth microglial gene among the RTL/SIRH genes [10] and that the RTL4 protein is secreted in the developing brain and its secretion is enhanced by various stimuli, possibly related to NA.

## Results

### RTL4 is expressed in the postnatal brain

*Rtl4*CV KI mice allowed the detection of RTL4CV protein expression in the postnatal brain: it was detected mainly in the midbrain, hypothalamus, amygdala and hippocampus, mostly as an extracellular secretory protein. To determine the location and dynamics of the RTL4 protein in the brain, we generated *Rtl4*CV KI mice (Fig. 1A and Fig. S1). Immunoaffinity experiments using anti-Venus antibody revealed that the RTL4CV protein was detected in the 2w brain at the expected molecular weight (61 kDa: RTL4 (34 kDa) + Venus (27 kDa)) (Fig. 1B).

**Figure 1.**
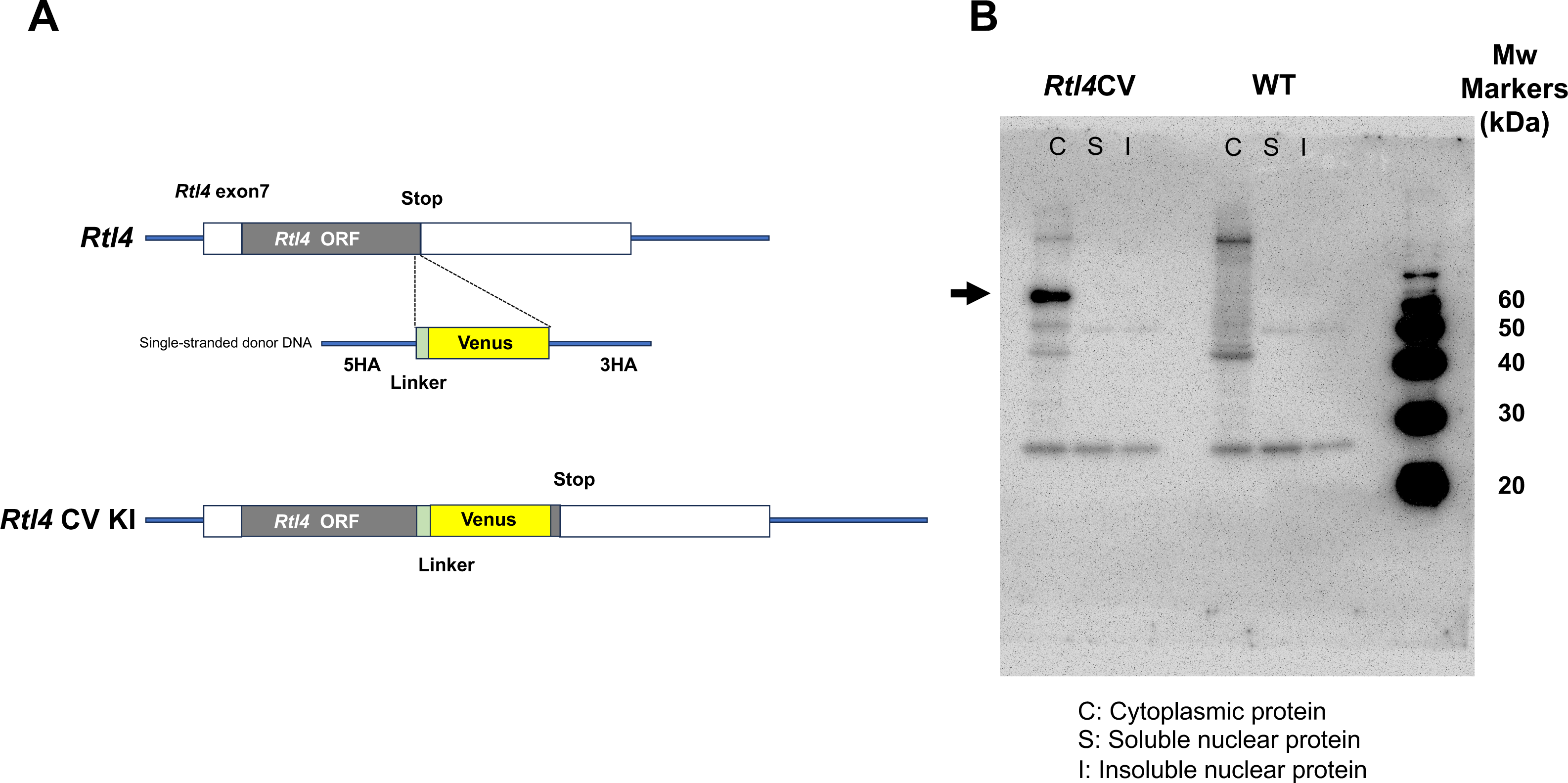
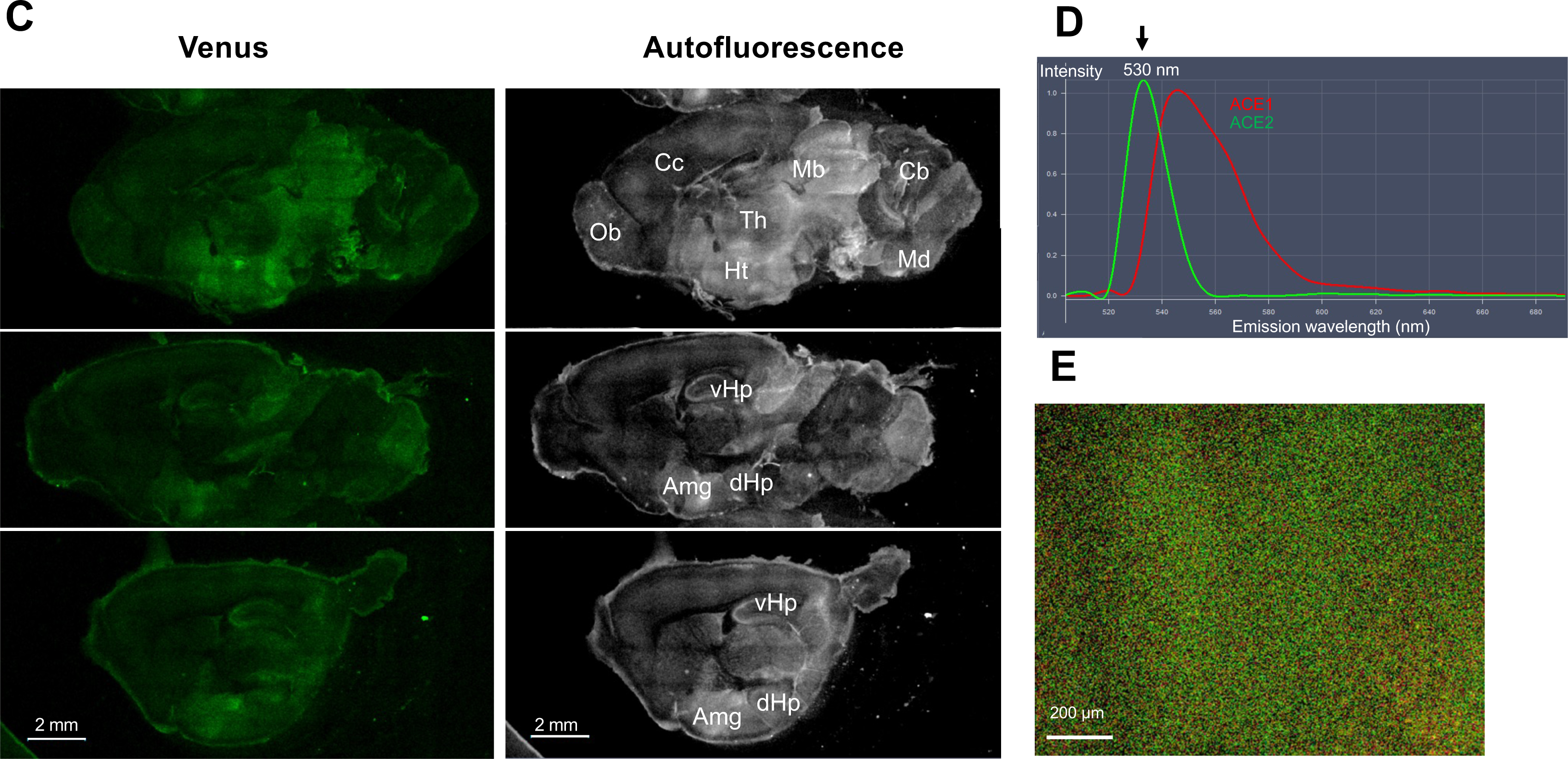
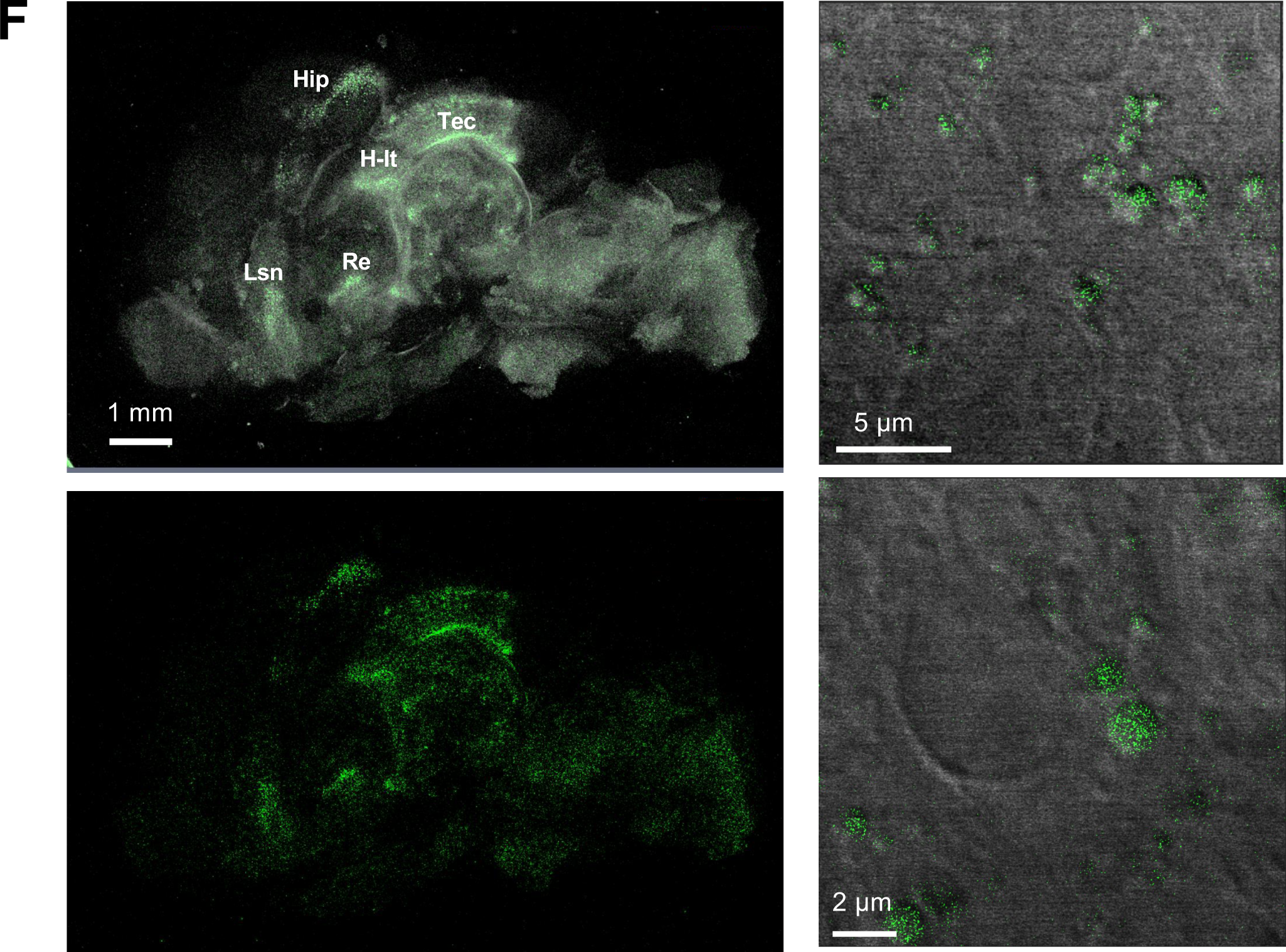
Detection of RTL4CV protein in the postnatal brain. **(A)** Construction of the *Rtl4* CV knock-in mouse. Schematic representation of wild type *Rtl4* and the modified genome structure of the *Rtl4* CV mouse. The mouse *Rtl4* open reading frame (gray box) is located on exon 7 (white box). The Venus coding sequence (yellow box) was inserted in front of the *Rtl4* stop codon together with a (GGS)x4 linker (light green box). See the details in Fig. S1. **(B)** Immunoaffinity experiment of the RTL4-CV protein in the 2 w brain. Immunoprecipitation was performed using an anti-GFP antibody. The estimated molecular weight of the RTL4CV protein is 61 kDa (RTL4 and Venus, 34 and 27 kDa, respectively). **(C)** Detection of RTL4 CV in P15 brain. Left: Venus fluorescence image. Right: Autofluorescence image. Amg: amygdala, Cb: cerebellum, Cc: cerebral cortex, dHP: dorsal hippocampus region, Ht: hypothalamus, Mb: midbrain, Md: medulla oblongata, Ob: olfactory bulb, Th: thalamus, vHp: ventral hippocampus region. **(D)** and **(E)** Venus signal detected by the LSM880 confocal laser scanning microscopy. The Venus signal (530 nm) was detected as the second strongest peak (D) in the hypothalamus (E). ACE: Automatic Composition Extraction. **(F)** Extracellular granules containing RTL4CV in P2 brain. Top left: Merged image of autofluorescence and Venus. Bottom left: Venus image. Right: Extracellular particles in the hippocampus. Merged images of transmission and Venus. Hip: Hippocampus, H-It: Habenulo-interpedincular tract, Lsn: lateral septal nucleus, Re: thalamus nucleus reuniens, Tec: Tectal commissure.

The RTL4CV Venus signal was confirmed by confocal laser scanning microscopy (Fig. 1C). The fluorescence from the RTL4CV protein was detected using a waveform with a peak at 530 nm emitted by Venus, calculated by the Automatic Composition Extraction (ACE) function (Figs. 1D and E). *Rtl4* mRNA expression levels were roughly estimated to be very low in the postnatal brain by qPCR (Fig. S2). The *Rtl4/β-actin* ratio increased around day 10 and peaked around 3w. This was more than 10-fold higher than the P1 level, but still less than 1/4,000 of the *β-acti*n level (Fig. S2). Although its mRNA level is very low in the P1 brain, Venus signals were detected in small but restricted regions, such as the midbrain and medulla oblongata (Fig. S3). In the P5 brain, a strong signal was detected in the midbrain, and a moderate signal was detected in the medulla oblongata and throughout the hypothalamus (Fig. S4). In the P9 and P10 brain, the hypothalamic signal increased to the same level as the midbrain (Fig. S4). After 2w, the hypothalamus became the main site of expression and the amygdala and hippocampus signals were also detected in the P15 brain (Figs. 1C and D). The hypothalamic signal increased until 3-4w, and after that it was maintained at a lower level than 4w, and the expression profiles remained the same. Most of the RTL4CV signal was detected as diffuse dots in the extracellular space (Fig. 1E), suggesting that it exists as a secretory protein. The exception is the early postnatal period, such as P1 and P2: signals were detected in the midbrain and medulla oblongata in the P1 brain (Fig. S3) and in the hippocampus, midbrain, lateral septal nucleus, thalamus nucleus reuniens and around the thalamus in the P2 brain (Fig. 1F, left), the majority of these signals were in the form of extracellular granules (1 µm in size) (Fig. 1F, right), suggesting that RTL4 may exist as a complex during this period. In the WT brain, no intrinsic Venus signal was detected within ACE10 in any postnatal development stage, so their signal intensities were calculated using the control Venus waveform and treated as experimental background (BG) (Figs. S5 and S6, see also Figs. 2A and B).

**Figure 2.**
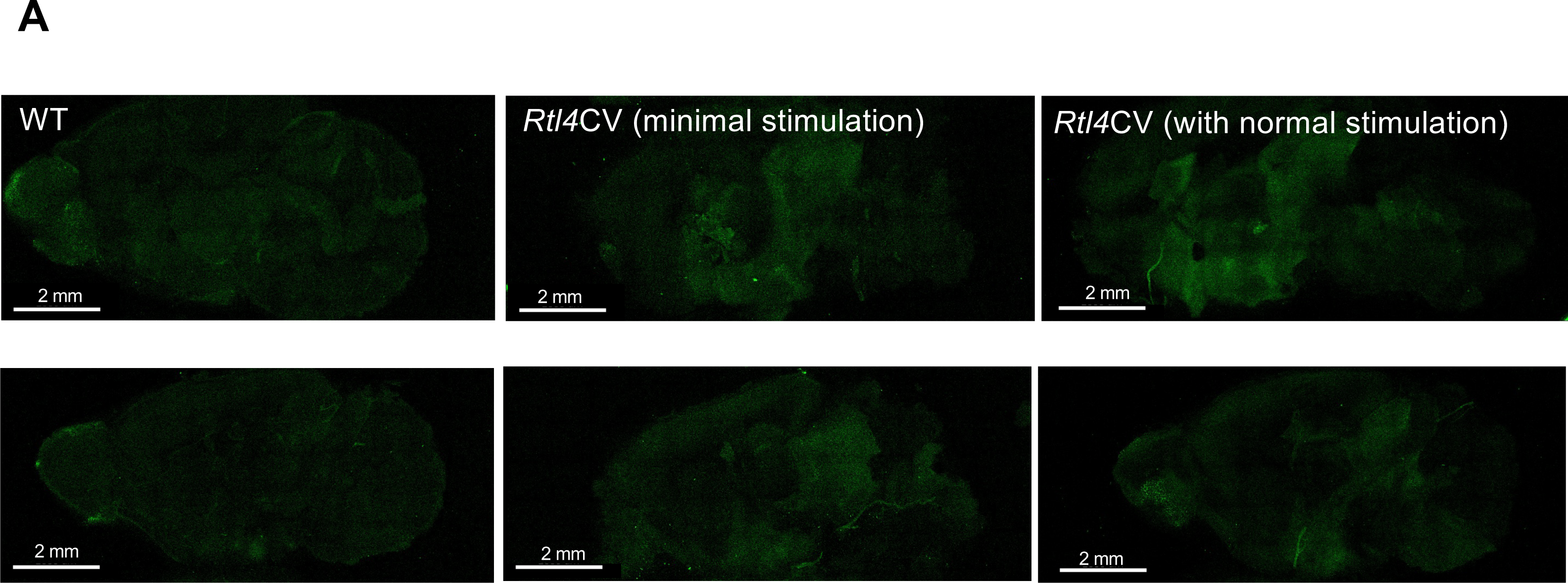
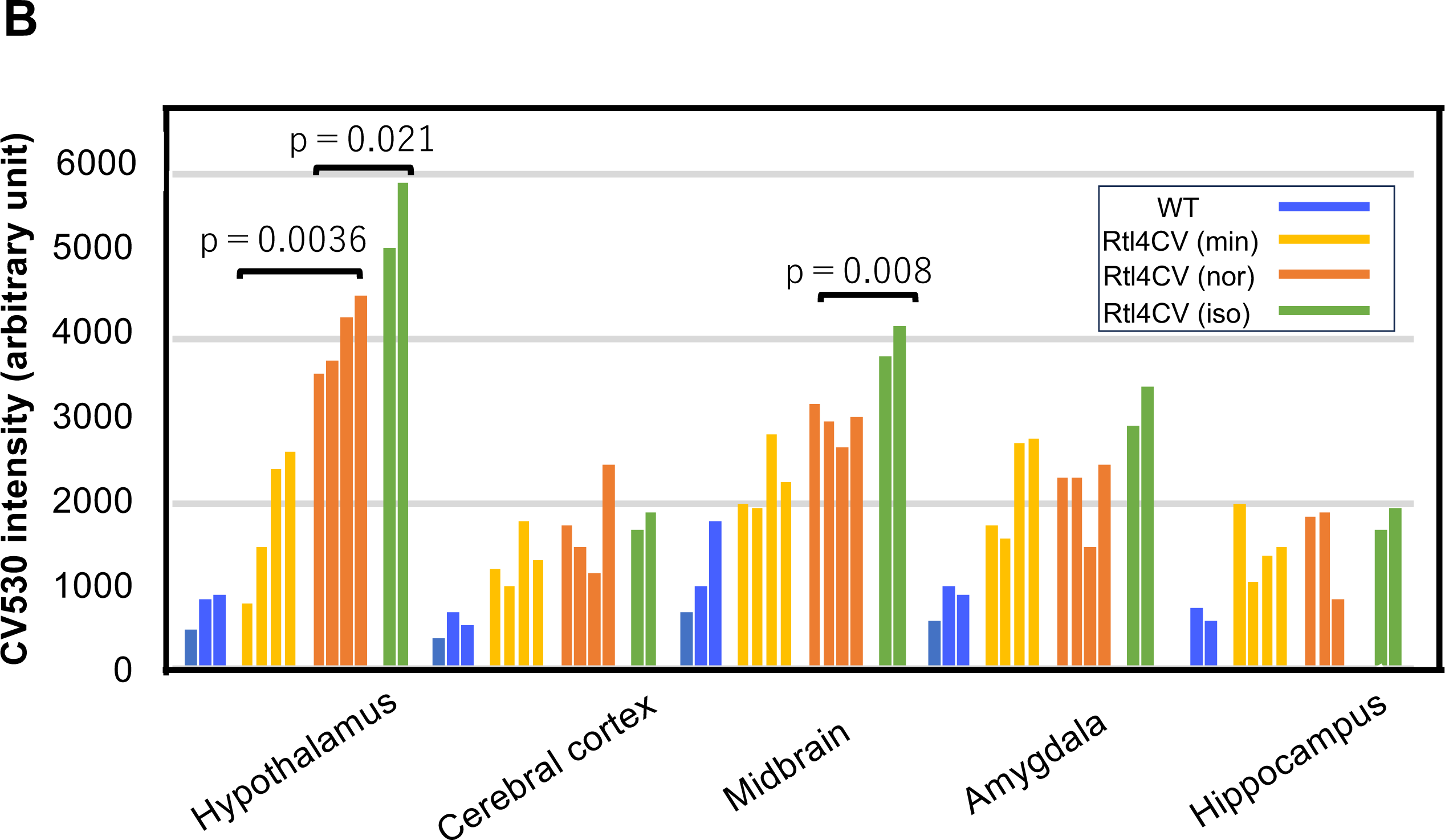
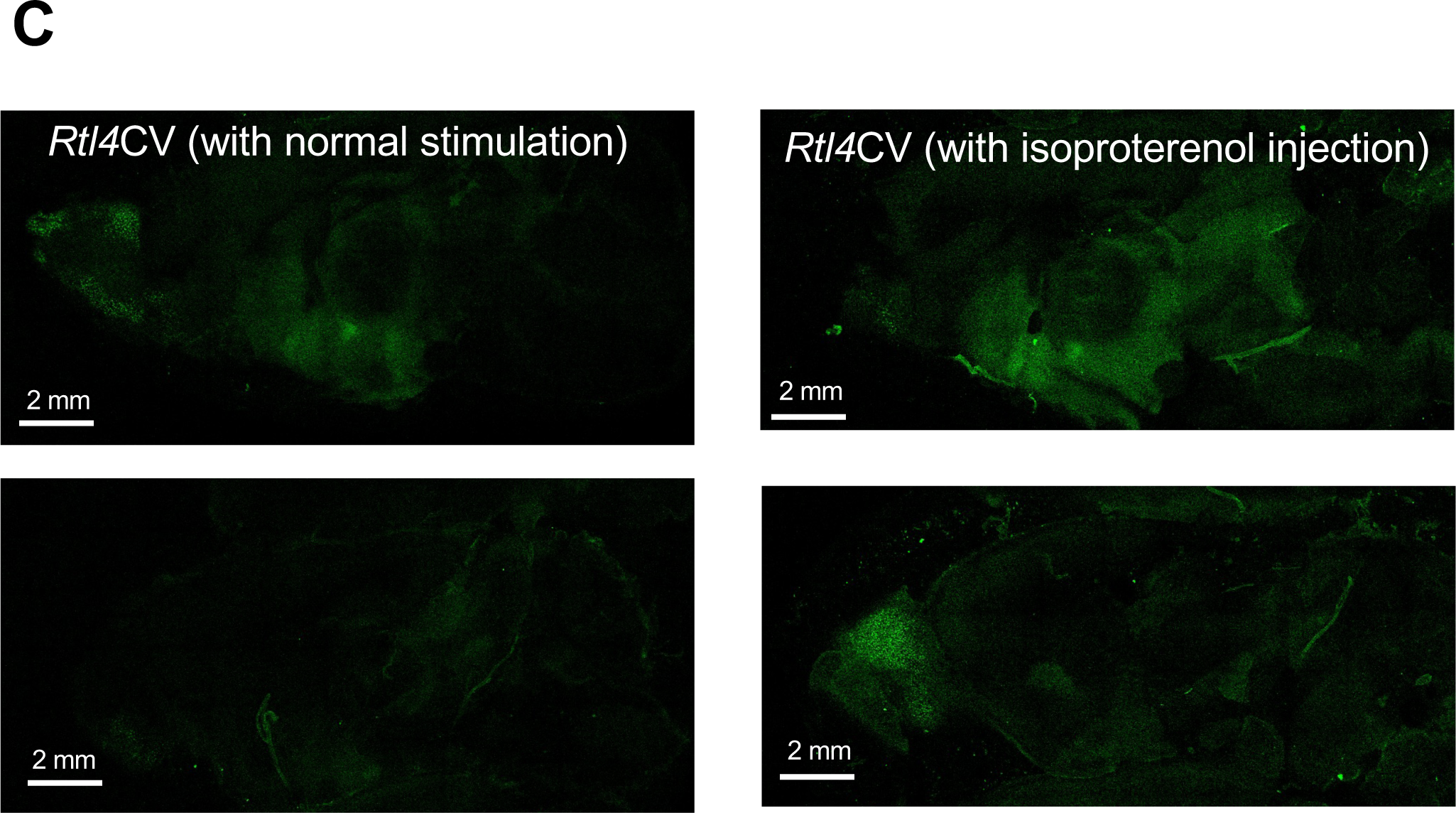
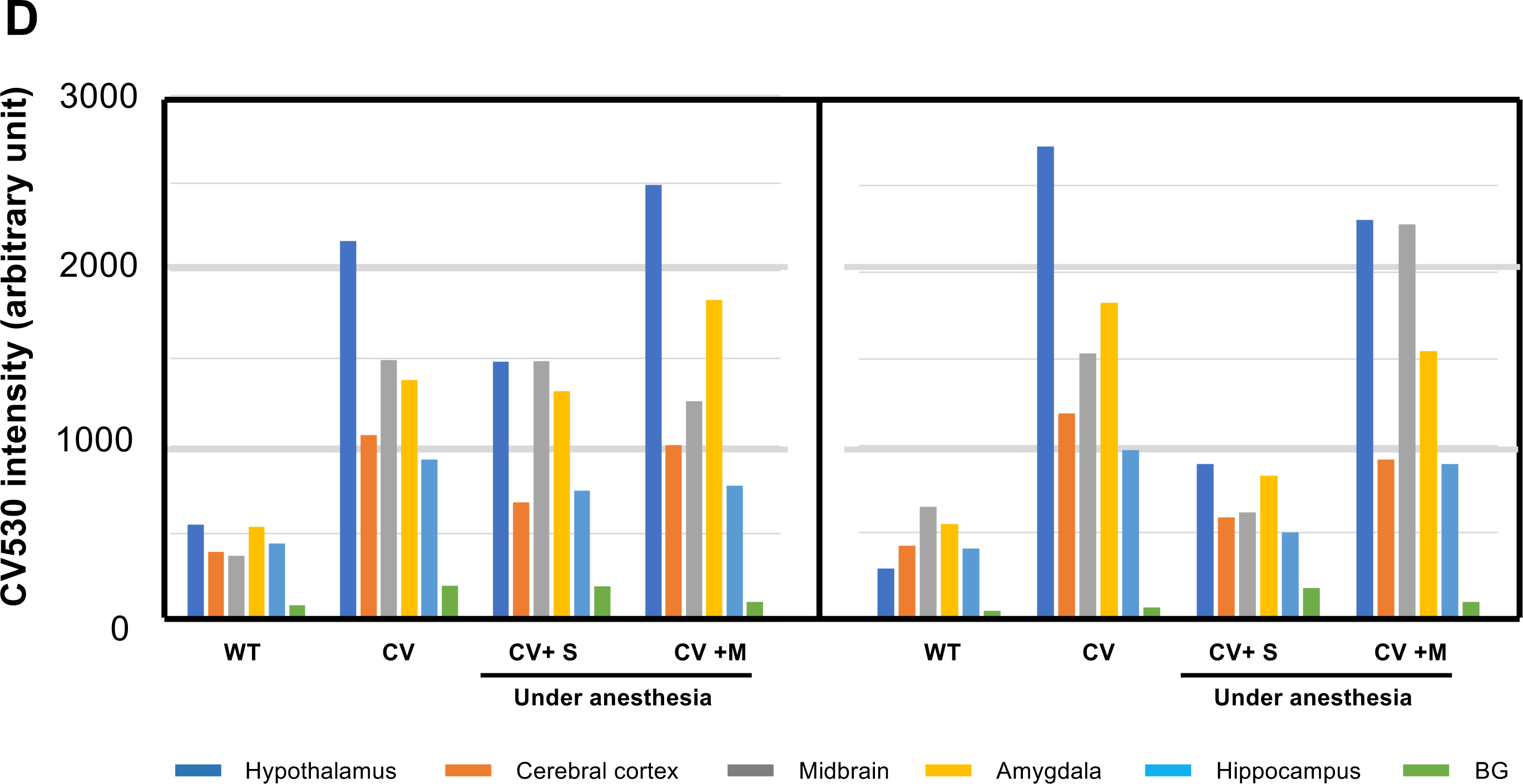
RTL4CV expression in the brain. **(A)** The effect of handling on RTL4CV protein expression in P21 brain. Venus images of WT (left), Rtl4CV with “the minimal stimulation” (middle) and Rtl4CV in “the normal condition” (right). Top: Inner side of the brain hemisphere. Bottom: Brain slice of 1.5 mm width. Bar: 2 mm. **(B)** Measurement of Venus signal intensity at different parts of the P21 brain (see also Fig. S7). Blue: WT, yellow: *Rtl4*CV with “the minimal stimulation (min)”, orange: *Rtl4*CV under “the normal conditions (nor)”. Green: *Rtl4*CV with isoproterenol cerebroventricular injection (iso, 10 μg/P21 mice). Each bar represents one P21 individual. The intensity of *Rtl4*CV in the hypothalamus under “the normal conditions” is significantly higher than that of *Rtl4*CV with “the minimal stimulation” (p=0.0036, two-tailed t-test). The intensity of *Rtl4*CV in the hypothalamus and midbrain is further increased after isoproterenol injection (p=0.021 and 0.008, respectively, two-tailed t-test). **(C)** Effect of isoproterenol on RTL4CV expression in P21 brain. Venus image of *Rtl4*CV under “the normal condition” (left) and *Rtl4*CV with isoproterenol injection. Top: Inner side of the brain hemisphere. Bottom: Brain slice of 1.5 mm width. Bar: 2 mm. **(D)** Effect of anesthesia and milnacipran on RTL4CV intensity. Left: WT (P28), *Rtl4*CV (P25) and *Rtl4*CV (P25) under isoflurane anesthesia. Right: WT (P30), *Rtl4*CV (P27), *Rtl4*CV (P27) with saline injection (CV+S) and *Rtl4*CV with milnacipran injection (CV+M). After the *Rtl4*CV mice were anesthetized with isoflurane for 1min, 10 μl of saline or milnacipran (total 167 μg) was administered intraventricularly, followed by additional anesthesia for 5 min. The signal intensity of hypothalamus (blue), cerebral cortex (orange), midbrain, (gray) amygdala (yellow), hippocampus (light blue) and background (light green) are presented.

### RTL4CV signal intensity varied in response to stress, environmental changes and the administration of isoproterenol and milnacipran

The patterns and intensities of RTL4CV expression varied in response to several factors related to NA. The previous *Rtl4* KO study suggested the relationship between RTL4 and NA. We hypothesize that RTL4CV is somehow responsive to NA, and this was supported by the finding that the intensity of RTL4CV signals was highly dependent on the experience of the *Rtl4*CV mice prior to observation. In the above analyses in Fig. 1C and Figs. S3-S4, the brain was observed after the mice were transferred from the breeding room to the laboratory (hereafter referred to as “the normal (or usual) condition”). However, the RTL4CV signal intensity of the hypothalamus and midbrain was significantly lower when the mice were treated in the breeding room before transfer (hereafter as “the minimal stress condition”) (Figs. 2A and B, Fig. S7), suggesting that these regions are highly sensitive to the various stimuli and/or stressors, such as differences in room brightness, ambient noise and vibration during cage transport. Even in the minimal stress condition, the signal intensity was variable, presumably due to inter-individual differences in sensitivity and/or slight differences in handling. Since NA plays an important role in arousal, attention, vigilance and stress responses, these results seem consistent with our hypothesis. An experiment with intraventricular administration of isoproterenol (10 μg per P21 brain) [24], an adrenergic receptor beta agonist that mimics the effects of NA, confirmed a significant increase (25%) in RTL4CV signal intensity in the hypothalamus and midbrain compared to those in “the normal condition” (Figs. 2B and C).

Furthermore, we also observed that isoflurane anesthesia of *Rtl4*CV mice consistently decreased signal intensity (Fig. 2D, left), whereas it was maintained by intraventricular administration of milnacipran (total 167 ng) [27] (Fig. 2D, right), suggesting that the RTL4CV signal intensity depends on the arousal state and is stabilized in the presence of NA due to the inhibition of the NA reuptake.

### RTL4CV is expressed in microglia and responds to isoproterenol

Cultured cell experiments confirmed that RTL4CV is expressed in microglia and that RTL4CV secretion is responsive to isoproterenol. Since the distribution of RTL4CV protein as a secretory protein is similar to that of RTL5 and RTL6 proteins secreted by microglia [36], we tested whether microglia express RTL4CV by performing mixed glial culture experiments on glia isolated from neonatal KI mouse brains (P0 or P1) [36, 37, 44]. Under this *in vitro* condition, several types of microglia, such as flat, spindle-shaped and floating round cells, were observed within, on and above astrocyte feeder cells, respectively, by the immunofluorescence staining with anti-Iba1 antibody (Fig. S8 and supplementary movie). The RTL4CV signal was detected in all of microglia types but not the astrocyte feeder cells (Figs. 3A and B). Interestingly, the RTL4CV signal intensity was also higher in the culture medium of *RTL4*CV than that of the control WT microglia, suggesting that RTL4CV is secreted into the culture medium under this condition (Fig. 3B). Although the possibility of RTL4CV expression in neurons and/or oligodendrocytes cannot be completely excluded, these results demonstrated that the RTL4C protein is expressed in microglia and secreted into the brain.

**Figure 3.**
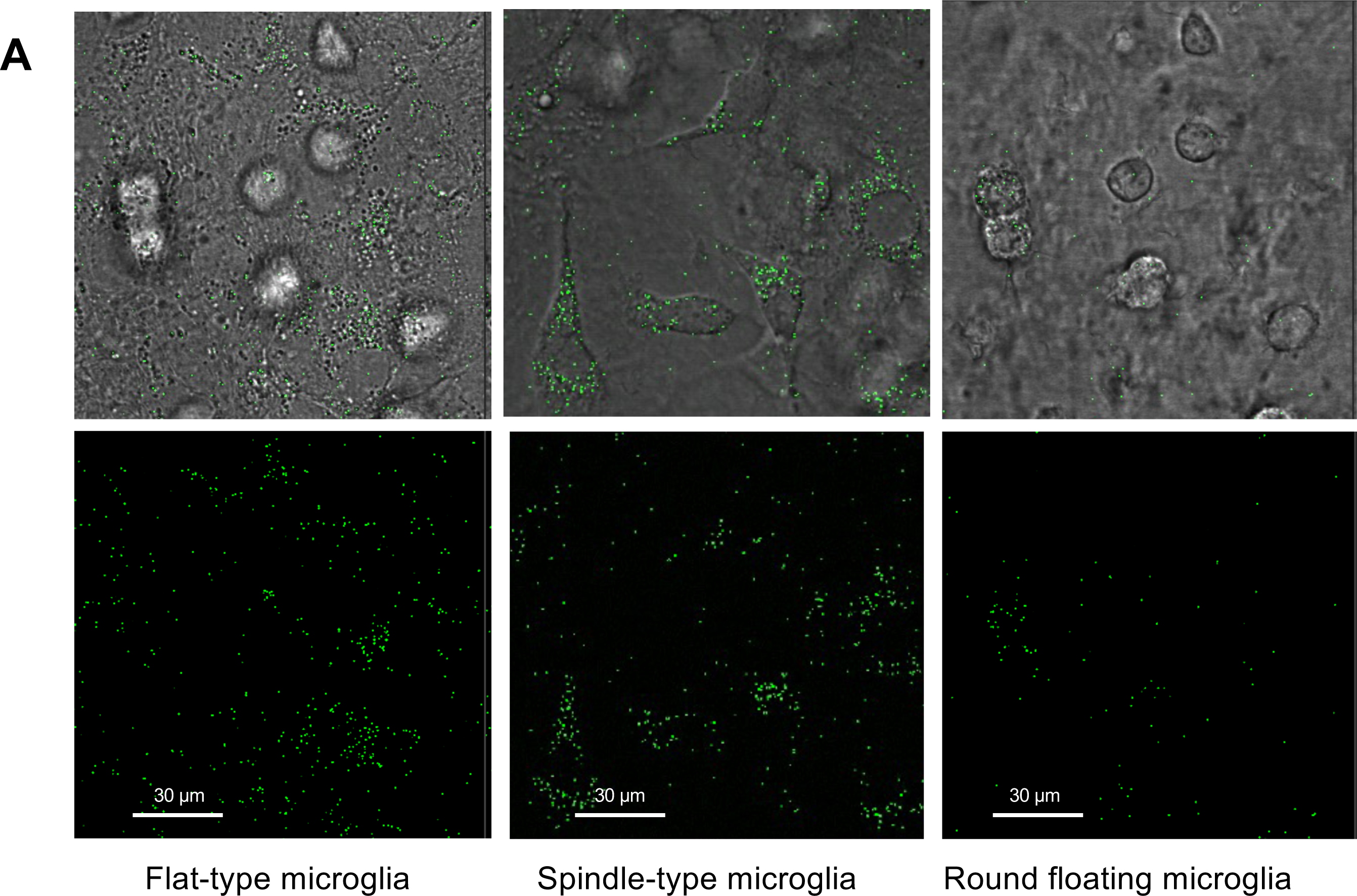
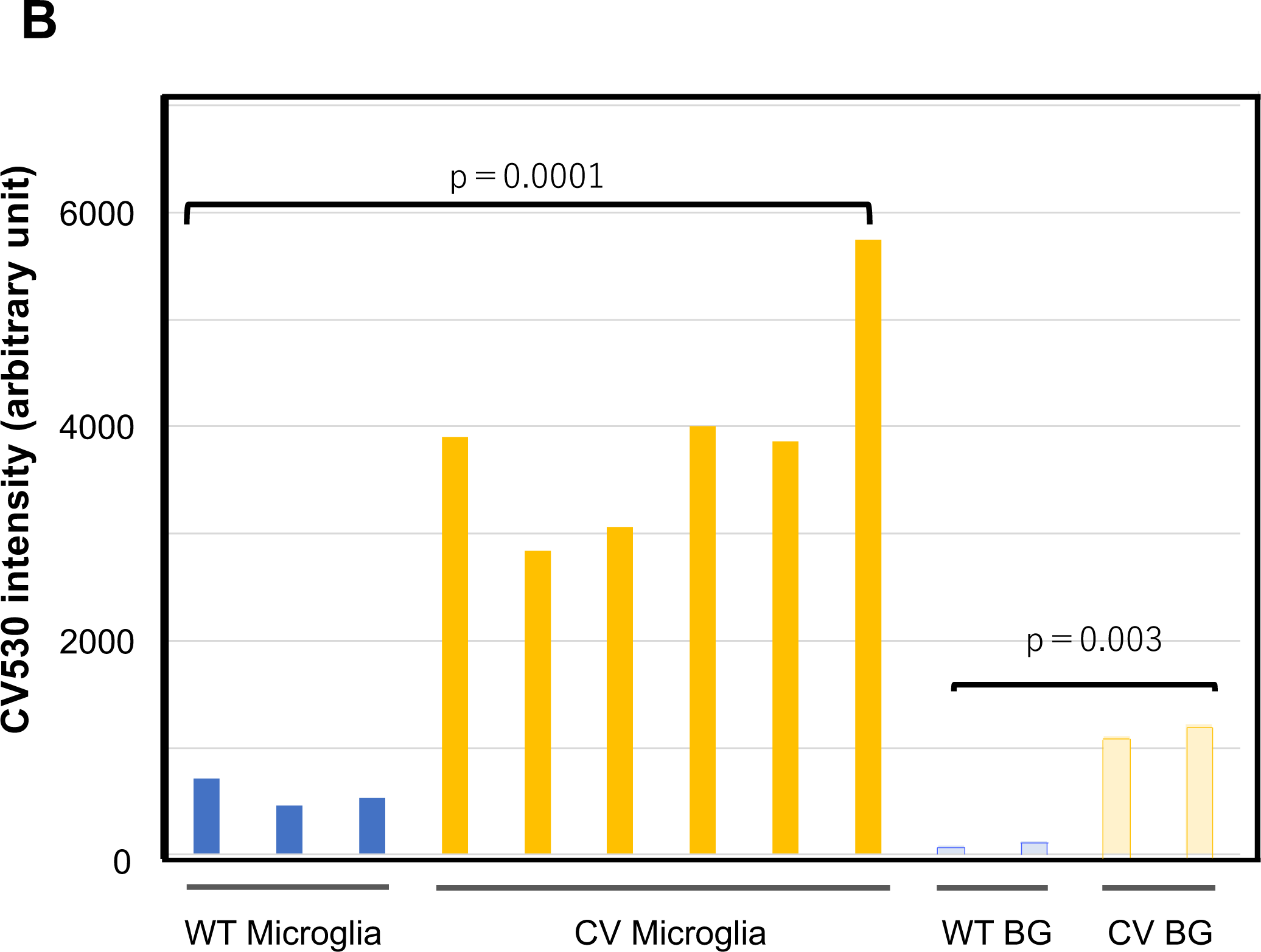
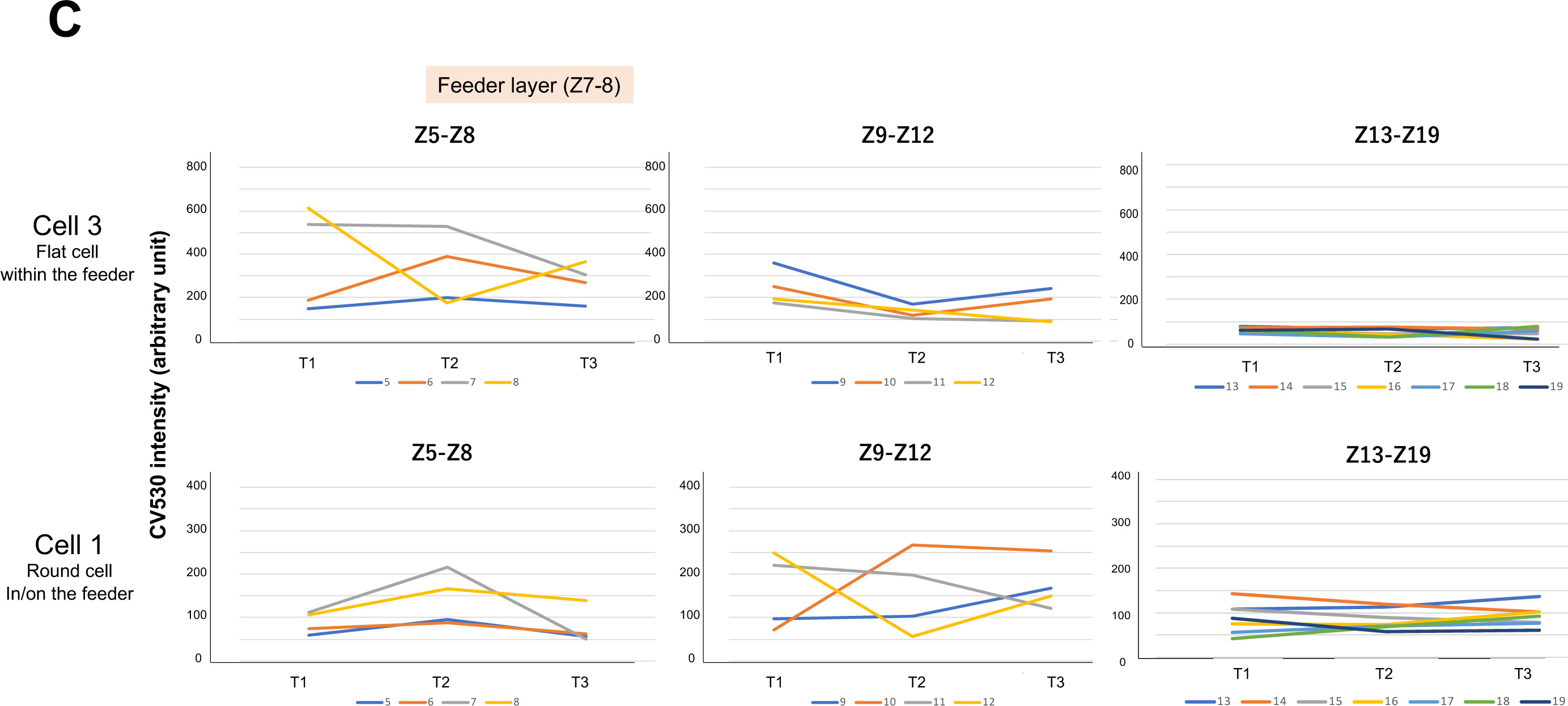
RTL4CV expression in cultured microglial cells. **(A)** Different types of microglia expressing RTL4CV in the mixed glial culture: (left) flat type, (middle) spindle type, (right) floating round type (see also Fig. S8). Top: Merged image of transmission and Venus. Bottom: Venus image. Bar: 30 μm. **(B)** Measurement of Venus signal intensity of microglial cells isolated from WT (blue) and *Rtl4*CV (yellow) (left) and that of culture media from WT (light blue) and Rtl4CV (pale yellow) (right). The intensity of *Rtl4*CV microglia is significantly higher than that of WT (p=0.0001, two-tailed t-test). The same is true for the culture media (p=0.003, two-tailed t-test). **(C)** Rapid secretion of RTL4CV in response to isoproterenol administration. Each of the 3D section intensity data (at 0.5 μm interval, total of 19 Z positions) from the mixed glial culture dishes was obtained in the time-lapse experiment (90 s interval). Isoproterenol (20 μM) was added to the culture media during the 90-s interval between T1 and T2. Therefore, the T2 data were obtained 30 s after administration. As shown in the top label, the astrocyte feeder layers were located at Z7-Z8, and the microglial cells were located from Z8 (within the feeder) to Z11-Z12 (on the feeder) according to their morphology. The signals at the Z position numbers, Z5-Z8, Z9-Z12 and Z13-Z19, are displayed. The results of the two microglial cells (No. 1 and 3 in Fig. S9) are shown. The results of Z1-Z4 (bottom side) are in Fig. S9.

Furthermore, rapid changes in signal intensity were observed in and around the microglia immediately after isoproterenol (20 μM) administration into the culture media [25]. We analyzed the results of the three-dimensional (3D) time-lapse experiments in the mixed glial culture before (T1) and after (T2 and T3) isoproterenol administration (Fig. 3C). The astrocyte feeder layers were located at Z7-Z8, and the microglial cells within the feeder were located approximately at Z8 and those on the feeder at Z11-Z12. The signal intensities of the microglial cells decreased after isoproterenol administration (e.g., at Z8 in Cell 3 and Z12 in Cell 1, whereas those of the area beneath the microglial cells increased in the opposite manner (e.g., at Z6 in Cell 3 and Z10 in Cell 1). The intensity of the other Z positions (e.g., Z1-4 and Z13-19 in Cell 3 and Cell 1) remained unchanged (Fig. 3C and Fig. S9). These results suggest that the microglial cells secreted RTL4CV into the media in response to isoproterenol.

## Discussion

### *RTL4,* a causative gene in ASD, is a microglial gene

*RTL4* has been implicated in ASD [14], and *Rtl4* KO mice exhibited behavioral abnormalities, such as increased impulsivity, reduced adaptation to novel environments and impaired short-term spatial memory as well as low recovery rate of NA in the frontal cortex [15]. In this study, we demonstrated that a causative ASD gene, *Rtl4,* is a microglial gene that responds to the NA signaling in the brain. *In vitro* experiments demonstrated that RTL4CV is expressed in microglia, not astrocytes, and also suggested that the RTL4CV secretion responds rapidly to isoproterenol administration. The dynamics of RTL4CV protein in the postnatal brain, such as its dependence on arousal state and its highly sensitive response to stress/environmental changes, including the response to isoproterenol and milnacipran, would provide crucial information in elucidating the etiology of ASD. As the NA response to stressors is likely to be reduced under anesthesia, RTL4 may also be a good indicator of arousal state and/or NA secretion in certain brain regions.

It has been shown that stress and the associated regulation of corticotropin-releasing factor (CRF) and opioid hormones affect the development of the NAergic system in the brain, and that this adaptive response to stress promotes behavioral flexibility [45–47]. Since RTL4 secretion is induced by various stresses, such as handling and transfer from the breeding room to the laboratory (Figs. 2A-D), the lack of this response from postnatal to adult life may explain why *Rtl4* KO mice exhibit reduced adaptability to a new environment [15], possibly due to impaired development of the NAergic system. The RTL4CV signal remained upon the administration of milnacipran (Fig. 2D), suggesting that the RTL4CV intensity is dependent on the presence of NA, although the mechanism by which the RTL4CV signal is stabilized by NA remains unclear. Is it possible that secreted RTL4 forms a complex with NA (see last section) or that secreted RTL4 is somehow involved in the reuptake of NA in neurons? Assuming that RTL4 has the ability to support the NA reuptake process, this may be consistent with the previous findings that *Rtl4* KO exhibited increased impulsivity and slow recovery of NA levels in the prefrontal cortex [15], because NAT is responsible for reuptake of most of the secreted NA back to the presynaptic noradrenergic neurons [26]. The resulting low level of NA in the NA neuron may be related to the impaired short-term spatial memory of *Rtl4* KO mice [15, 48]. Taken together, RTL4 is a novel and important microglial factor in the brain response to NA and may serve as an important therapeutic target in ASD, although further studies are needed to elucidate the exact function of the secreted RTL4 protein in the NA reuptake process.

### Importance of RTL4 in the developing brain and its relationship to the LC-NA system

What is the role of RTL4 secretion the brain? Since NA reportedly shapes the properties of NAergic neuronal networks in the developing brain [28, 29], it is likely that the impaired NA response during the postnatal period interferes with the normal development of NAergic neuronal networks. Importantly, the timing and location of RTL4 expression as well as its induction closely correspond to critical periods and regions of brain development. Learning to recognize the maternal odor is critical for the survival of infants in order to receive proper maternal care, known as infant attachment learning. Incidentally, β-ARs play an important role in infant attachment learning, and the absence of α2-AR activity during this period is also considered important [29]. During this early postnatal period, RTL4CV was present as mostly 1 µm extracellular granules in the hippocampus, midbrain, lateral septal nucleus, thalamus nucleus reuniens and around the thalamus (Fig. 1F). The hippocampus is the is well-known center of memory and the lateral septal nucleus, with its abundant inputs from neocortical and allocortical regions, is ideally positioned to integrate perceptual and experiential signals and plays a critical role in emotionality, social behavior and feeding processes, through neural connections with the hippocampus, midbrain, thalamus nucleus reuniens and hypothalamus [49, 50]. It is therefore highly likely that RTL4 plays an important role in this process.

The intrinsic RTL4CV signal was very strong around P10 (Fig. 1C). After P10 in rats, neonates acquire aversion and fear learning [29]. The inhibitory effect of α2-AR is activated subsequent to P10 in rats and is known to play an important role in the subsequent formation of the NAergic neuronal network. The interaction between LC activity and NA signaling in the hypothalamus and hippocampus, as well as corresponding activity in the amygdala, has been suggested to be important for fear learning in early development. Since amygdala expression was observed in the P10 mouse brain under “the normal conditions” (Fig. S11), it is possible that RTL4 also takes part in this process.

The RTL4CV signal peaks at 2-4 weeks (2-4 w) (Fig. 1C and Fig. 2A). In rats, robust synaptogenesis in the noradrenergic pathway occurs between P10-15 and P20-30. α-ARs are present in the newborn, but their number increases at most sites after birth, peaking around P15 for α2-ARs [51] and between P15 and 20 for α1-ARs [29, 52]. In the rat cerebral cortex, β-AR density increases gradually from P5, reaching higher than adult levels by 2-3 w and then maintaining adult levels for several months [29]. This is in good agreement with RTL4 profiles in these mice (Figs. 1C and 2A, Fig. S2). It is likely that various different stimuli are able to induce RTL4 expression in different brain regions so as to somehow promote the new NAergic neuronal network (Figs. 2A-D).

### *Rtl4* is the fourth RTL/SIRH gene identified in eutherian microglia

RTL/SIRH are genes derived from *Metaviridae*, retrovirus-like retrotransposons, by three independent domestication events, resulting in therian-specific *PEG10,* eutherian-specific *RTL1* (aka *PEG11*) and *RTL3-9* (*SIRH3-11*), respectively [1–11]. They all encode Gag-like proteins, but *PEG10*, *RTL1* and *RTL3* (aka *SIRH9*) also encode a portion of Pol-like protein. They are good examples of exaptation proposed by Gould and his colleagues [53, 54], because each of the ten RTL/SIRH gene has a unique, essential and/or important function in the placenta and/or brain [8, 10]. *PEG10*, *RTL1* and *LDOC1* (aka *RTL7* or *SIRH7*) play both essential/important roles in the placenta [5, 55–59] and in the brain as neuronal genes along with *RTL8A, B, C* (aka *SIRH5, 6, 4*) [60–65], whereas *RTL5* (aka *SIRH8*), *RTL6* (aka *SIRH3*) and *RTL9* (aka *SIRH10*) play roles in the innate immune system as microglial genes in the brain against viruses, bacteria and fungi by removing dsRNA/ssDNA, LPS and zymosan, respectively [36, 37].

Thus, *RTL4* is the fourth microglial gene among the RTL/SIRH genes [10], suggesting that these four RTL genes contribute to the eutherian-specific microglia. What is the role of RTL4 in microglia? Apparently, RTL4 responds to psychological stressors via NA, whereas RTL5, RTL6 and RTL9 respond to physiological stressors, i.e., various pathogens, suggesting that each of these RTL genes plays a distinct role in the brain’s stress responses in a eutherian-specific manner. The retroviral GAG from which each RTL/SIRH gene is derived encodes molecules that can self-assemble. RTL5 and RTL6 form large extracellular particles that bind specific PAMPs, as if taking advantage of this property of GAG. One possibility is that RTL4 supports the NA reuptake response, as discussed above, by forming a sort of RTL4-NA complex similar to the RTL5-dsDNA and RTL6-LPS complexes.

In the early postnatal period, similar to RTL5 and RTL6 [34], RTL4 was present as 1 µm extracellular granules in specific regions where new neuronal networks are thought to form (Fig. 1F, right) [49, 50]. Although it remains unclear what components are contained in these granules, it is interesting to speculate that RTL4 recognizes some damage-associated molecular patterns (DAMPs) and acts as a scavenger of these substances to maintain a healthy brain environment during this period, because it is possible that cellular debris was generated during NA-induced network remodeling [38, 66, 67]. This situation is similar to recognition receptors (PRRs), such as TLRs and cytoplasmic NOD-like receptors (NLRs), which recognize both PAMPs and DAMPs to orchestrate an inflammatory response [68, 69]. Regardless of its relationship to NA *per se*, it seems likely that eutherians have improved their use of NA with the acquisition of *RTL4* and that its defect leads to the development of ASD.

As microglia recently attracted considerable attention as an important factor in the pathogenesis of not only ASD but also other neurodevelopmental and neuropsychiatric disorders [38–41, 67, 70–72], further experiments are needed to elucidate the biochemical function of RTL4, as well as other, as yet unidentified microglia-specific genes, in the etiology of ASD. This study provides evidence not only for the role of novel microglial genes in ASD, but also for the unexpected contribution of a retrovirus-derived acquired gene to the evolution of the eutherian brain, together with other neuronal genes, such as *PEG10, PEG11/RTL1* and *RTL8A-C* [60–65], and microglial genes, such as *RTL5, RTL6 and RTL9* [36, 37].

## Materials and Methods

### Mice

All of the animal experiments were reviewed and approved by the Institutional Animal Care and Use Committees of Tokai University and Tokyo Medical and Dental University (TMDU), and were performed in accordance with the Guidelines for the Care and Use of Laboratory Animals of Tokai University and TMDU.

### Generation of the *Rtl4*CV mice

*Rtl4*CV mice were generated as previously described with minor modifications [34]. Genomic DNA near the stop codon of the *Rtl4* gene was cleaved using Cas9 and a guide RNA targeting 5’-AACGTTCTAGAACTCCAGCA-3’. A plasmid vector was constructed to introduce DNA encoding most of the C-terminal amino acids of *Rtl4* located downstream of the Cas9 cleavage site, a 12 amino acid linker ((GGAGGATCA)x4) and the Venus protein into the Cas9 cleavage site upstream of the *Rtl4* stop codon so that RTL4 and Venus were expressed as a fusion protein connected with the linker. The vector also had homology arms of 1500 bases at the 5’ and 3’ ends of the introduced sequence, and the codon encoding arginine 301 was mutated from AGG to AGA to prevent re-cleavage by Cas9 after genetic recombination. The single-stranded DNA was synthesized as a mutation donor as previously described [34] and used to generate the genetically modified mice. Briefly, double-stranded DNA was amplified using the primer pair (5’ to 3’) TGCGTCCACTACCAAAGGAT and pre-phosphorylated GAGGAGGGGCTATCTTTCAAAC, and its 5’-phosphorylated strand was digested with lambda exonuclease.

Founder mice were genotyped using the three primer pairs (5’ to 3’) of CCAGATTTGATCACTCAGTGC/TGTTGTGGCGGATCTTGAAG, AGCAGCACGACTTCTTCAAG/CTCTTGAAGCTGATTGGTCC and CCAGATTTGATCACTCAGTGC/CTCTTGAAGCTGATTGGTCC. Two founder mice had unexpected mutations in the 3’UTR of *Rtl4,* a 12 base pair insertion and a 1 base pair deletion in founder #10 and a 2 base pair deletion in #13. These founders were used to generate 2 *Rtl4*CV lines and their progeny were used for further experiments.

### Immunoprecipitation and Western blotting

Postnatal brain (2w) was powderized in liquid N_2_ using a Multi-beads shocker (MB1050, YASUI KIKAI, Osaka), and the “Cytoplasmic Fraction”, “Soluble Nuclear Fraction” and “Insoluble Nuclear Fraction” were obtained using the LysoPure^TM^ Nuclear and Cytoplasmic Extraction Kit (FUJIFILM, Code No. 295-73901). The powder samples of WT and *Rtl4*CV brains (86.7 mg and 67.8 mg, respectively) were dissolved in 1,000 µl of Nuclear Fractionation Buffer on ice for 10 min. After centrifugation (500 x g, at 4°C) twice for 10 min, the supernatants were mixed and used as the “Cytoplasmic Fraction”. The pellets were resuspended in 500 µl of Nuclear Fractionation Buffer and mixed vigorously. After centrifugation (500 x g, at 4°C) for 10 min, the supernatant was discarded and the pellet was resuspended in 500 µl of Nuclear Extraction Buffer on ice for 30 min with vigorous mixing at 10 min intervals. After centrifugation (20,000 x g, at 4°C) for 10 min, the resulting supernatant was used as the “Soluble Nuclear Fraction”. The pellet was resuspended in 500 µl of Nuclear Extraction Buffer with vigorous mixing. After centrifugation (20,000 x g, at 4°C) for 10 min, the supernatant was discarded and the pellet was resuspended in 500 µl of Nuclear Extraction Buffer with vigorous mixing. After centrifugation (20,000 x g, at 4°C) for 10 min, the supernatant was discarded and the pellet was resuspended in 400 µl of SDS Lysis Buffer with vigorous mixing and sonicated until the pellet was no longer visible. After centrifugation (20,000 x g, at 4°C) for 10 min, the supernatant was used as the “Insoluble Nuclear Fraction”.

Each supernatant was mixed with 20 µl of anti-GFP (RatIgG2a), Monoclonal (GF090R), CC, Agarose Conjugate (NACALAI TESQUE) and incubated overnight at 4°C. The agarose beads were washed four times with 500 µl of RinseBuffer, 50 mM Tris-HCl (pH 8.0) and 150 mM NaCl at 4°C. Then the beads were incubated with 30 µl of SDS sample buffer and incubated at 95°C for 5 min. The samples were applied to gel electrophoresis using a 10% acrylamide gel. Western blot analysis was performed using a standard protocol. After blotting on a Hybond-P (GE Healthcare) membrane, the RTL4CV protein was detected with an ECL Prime Western Blotting Detection kit (GE Healthcare) using an anti-GFP antibody (MBL, Code No. 598) and an anti-Rabbit Goat Immuno-globulins/HRP (DAKO, P0160) as the 1^st^ and 2^nd^ antibodies. Signals were detected with an AE-9300 Ez CaptureMG (ATTO, Tokyo).

### Imaging using Confocal Laser Scanning Fluorescence Microscopy

Fresh brain and brain slices (1.5 to 2 mm in depth) from *Rtl4*CV KI mice were used for analysis with a ZEISS LSM880 (ZEISS, Germany) without fixation. The samples were observed using a Plan-Apochromat lens (10x, numerical aperture =0.45, M27, ZEISS) and a C-Apochromat lens (63x numerical aperture =1.2 Water, ZEISS). The tiling with lambda-mode images was obtained using the following settings: pixel dwell: 1.54 μs; average: line 4; master gain: ChS: 1250, ChD: 542; pinhole size: 33 µm; filter 500 – 696 nm; beam splitter: MBS 458/514; lasers: 514 nm (Argon 514), 0.90 %. For the tiling-scan observations, the tiling images were captured as tiles: 84, overlap in percentage: 10.0, tiling mode: rectangular grid, in size: x; 15442.39 µm, y; 9065.95 µm. Spectral unmixing and processing of the resulting images was performed using ZEN imaging software (version 2.3 SPI, Carl Zeiss Microscopy, Jena Germany). The spectrum of the Venus proteins (Maximum peak emission fluorescence wavelength: 528 nm) was detected only in the *Rtl4*CV samples and not in the wild type control samples. ZEN Blue edition ver. 3.9 (ZEISS) was used to measure and correct the average intensity of fluorescent molecules and the average tissue and extracellular steady-state background fluorescence intensity from 2D, 3D and 4D tissue and cell fluorescence images. Briefly, the raw images were averaged to remove noise, and each component image (RTl4CV530, AF530, etc.) was acquired by spectral analysis. The target region areas (the cortex, hypothalamus, amygdala, midbrain, hippocampus, etc.) of the acquired images were manually identified for validity and only the target areas were analyzed. Evaluation images were acquired using IMARIS ver. 6.1 (BitPlane).

The 3D time-lapse images were obtained under the following conditions. Microglial cells were cultured in 35 mm glass bottom dishes in a humidified incubator with 5 % CO2 at 37C using the Signal-top incubation system S1 (ZEISS, Germany). Time series: 90 sec interval, 11 frames. After obtaining the first frame of the mixed glial culture, isoproterenol (20 μM) was added to the culture media during the 90-s interval, and the second to eleventh frames were obtained. The first 3 frames (first frame: untreated, second and third frames: after drug administration) are shown (Fig. 3C and Fig. S9). Z-stack: 0.5 μm steps 28 slices (13.713 μm). Scaling (per pixel): 0.26 μm x 0.26 μm x 0.51 μm. Image size (pixels): 512 x 512. Bit depth: 16 bit. Objective lens: C-Apochromat 63 x/1.2 W Korr M27. Filters: 499-695. Beam splitter: Lambda. MBS: MBS458/514. MBS-InVis: Plate. DBS1: Mirror. Laser: Ar 514 nm, 0.9 %. Scan mode: Line sequential. Pixel time: 1.54 μs. Line time 30.00 μs. Frame time 1.00 s.

### Primary mixed glial culture

Primary mixed glial cultures were prepared according to the protocol of Lian *et al.* with modifications [42]. Briefly, P0-P2 brains were homogenized in phosphate buffer saline (PBS) supplemented with trypsin and DNase (final concentration 0.05% and 0.1 mg/ml, respectively) by repeated pipetting 10 times and incubated in a 37°C water bath for 5 min, and this process was repeated 3 times. After the addition of Dulbecco’s modified Eagle medium (DMEM) (Thermo Fisher Science, Gibco^TM^ Catalog No.11995065) along with 10 % fetal bovine serum (FBS) (Thermo Fisher Science, Catalog No.10437028), the debris was removed through a cell strainer (EASYstrainer, mesh size:100μm, Greiner, Cat. No 542000). After centrifugation (400 x g, at 4°C) for 5min, the supernatants were aspirated and the pellet was resuspended with 5 ml warmed culture medium. After determining the cell density, the mixed glial cells were plated into 35 mm glass base dishes (IWAKI, Code No. 3910-035) or 25 cm^2^ flasks (Canted Neck, Corning, Cat. No. 430639) and incubated in a CO2 incubator with 5 % CO2, 100% humidity at 37°C. Media were changed the next day and every 3-4 days thereafter.

### Isoproterenol and milnacipran administration

Isoproterenol hydrochloride (FUJI FILM, Code No. 553-69841) was dissolved in 0.9% sodium chloride saline at a concentration of 1 mg/ml just before the experiment. Based on the previous report by Yabuuchi et al [24], a 10-μl aliquot (10 μg/P21 mice) was administered intracerebroventricularly with an 18G needle syringe, or 20 μM (final concentration) isoproterenol was added to the culture medium according to the previous report by Tomozawa et al [25]. Milnacipran hydrochloride (Tokyo Chemical Industry (TCI), Code No. M2133) was dissolved in 0.9% sodium chloride saline at a concentration of 50 mg/ml and stored at -80°C. A 10-μl aliquot (167 ng) was administered intracerebroventricularly with an 18G needle syringe according to the previous report by Bagchi et al [27]. Mice were anesthetized using an isoflurane anesthesia machine SN-487-OT (SINANO MFG Co Ltd, Tokyo) under the condition of 4% isoflurane for 1 min, then, saline or milnacipran was injected intracerebroventricularly under 2.5 % isoflurane and further anesthetized for 5 min.

## Supplementary Materials

Fig. S1: Construction of the *Rtl4* CV knock-in mouse; Fig.S2: Postnatal *Rtl4* mRNA expression in WT brain; Fig. S3: Expression of RTL4CV protein in P1 brain; Fig. S4: RTL4CV expression in P5-9 brains; Fig. S5: WT P21 brain analyzed by the Automatic Component Extraction (ACE) function; Fig. S6: WT P21 brain; Fig. S7: Environmental effect on the expression of RTL4CV protein in P21 brain; Fig. S8: Microglial cells in the primary mixed glial culture (Iba1 staining). Fig. S9: Time-lapse experiment using mixed glial culture (isoproterenol administration). Fig. S10: RTL4 expression in the amygdala in the P10 brain. Supplementary movie: Iba1 staining of mixed glial culture.

## Acknowledgements

The authors thank Takako Usami (TMDU) for technical assistance in the embryo transfer experiment to generate *Rtl4*CV KI mouse and Naoko Takayasu at Tokai University and Tokai University Support Center for Medical Research and Education for qPCR experiments and breeding the mice. Pacific Edit reviewed the manuscript prior to submission.

## Author Contributions

F.I., J.I., M.I., A.M., M. Y. and T.K.-I. performed the experiments and analyzed the data. A.M., T.S. and Y.H. generated *Rtl4*CV KI mouse. F.I. and T.K.-I. designed the study and wrote the manuscript. All authors agree to be accountable for the content of the work.

## Informed Consent Statement

Not applicable.”

## Data Availability Statement

Data can be available upon request from the corresponding author. *Rtl4*CV mice are available from RIKEN BioResource Center (Tsukuba, Japan) along with *Rtl4* KO, *Rtl5*CmCherry (CmC), *Rtl5* KO, *Rtl6*CV, *Rtl6* KO, *Rtl9*CmC and *Rtl9* KO mice.

---

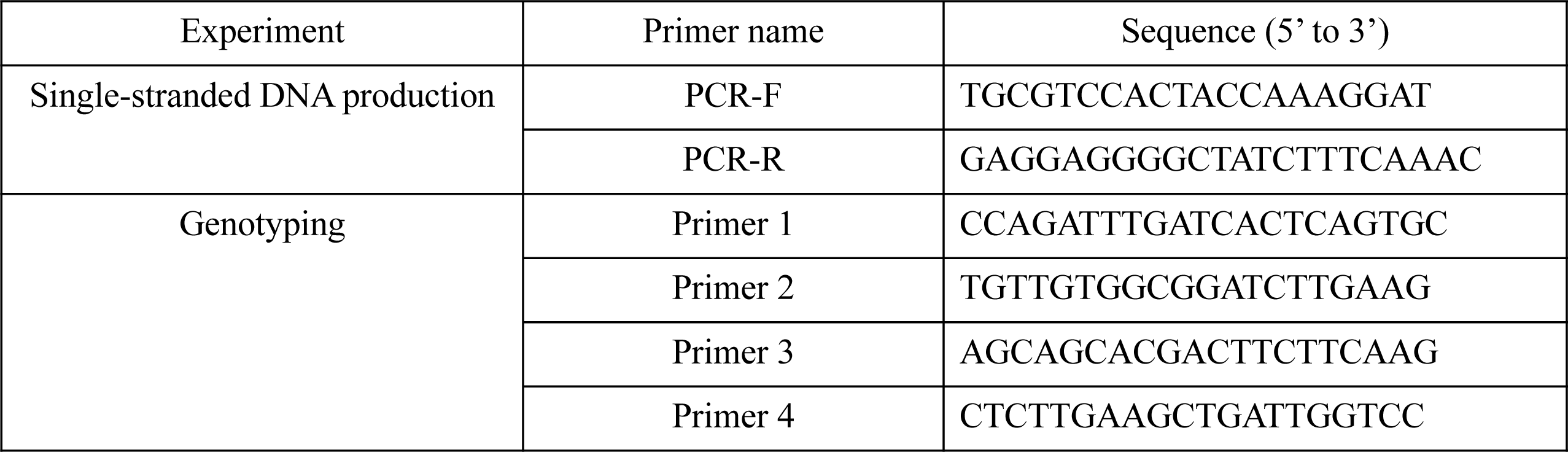
Primers used for *Rtl4CV* mouse generation.

**Fig. S1.**
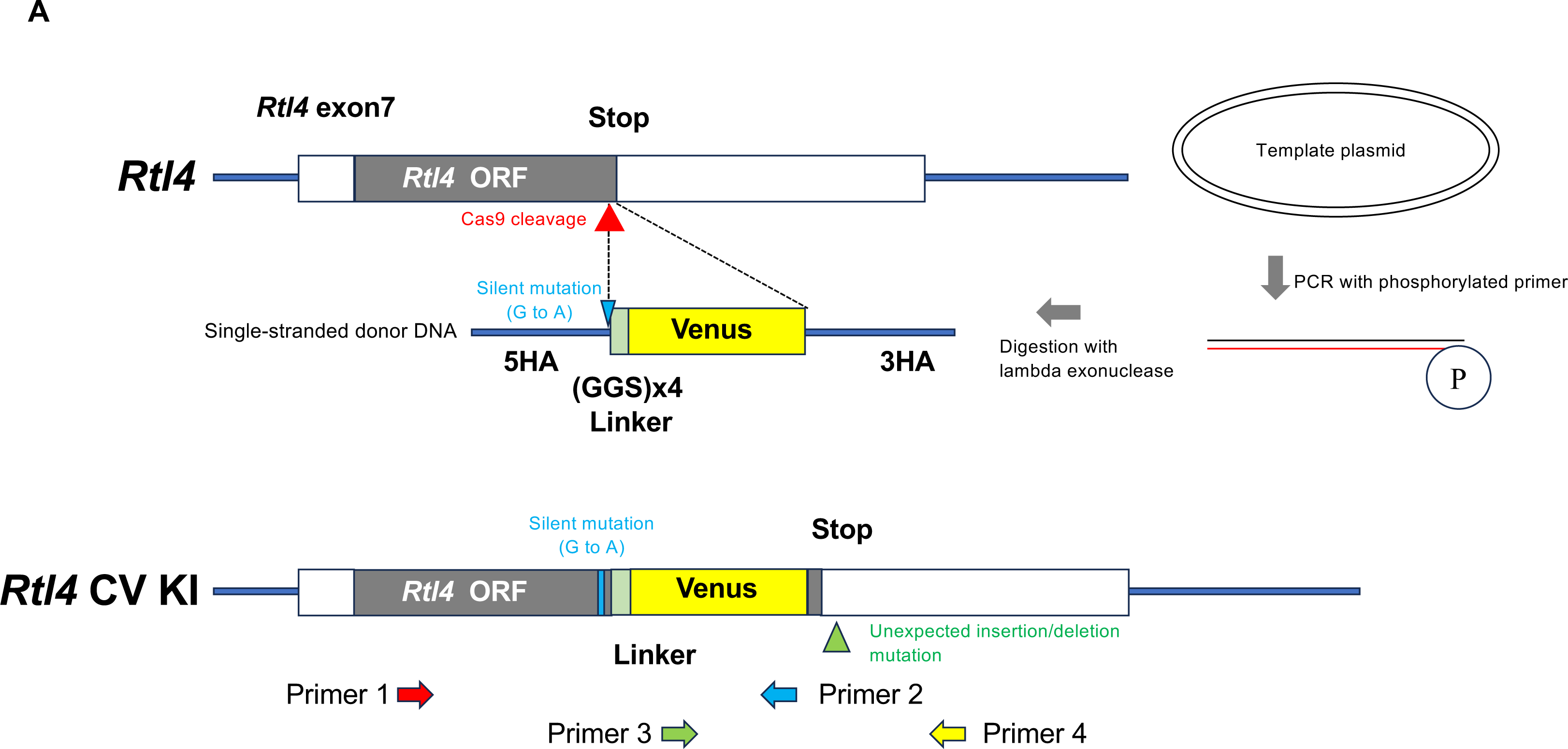
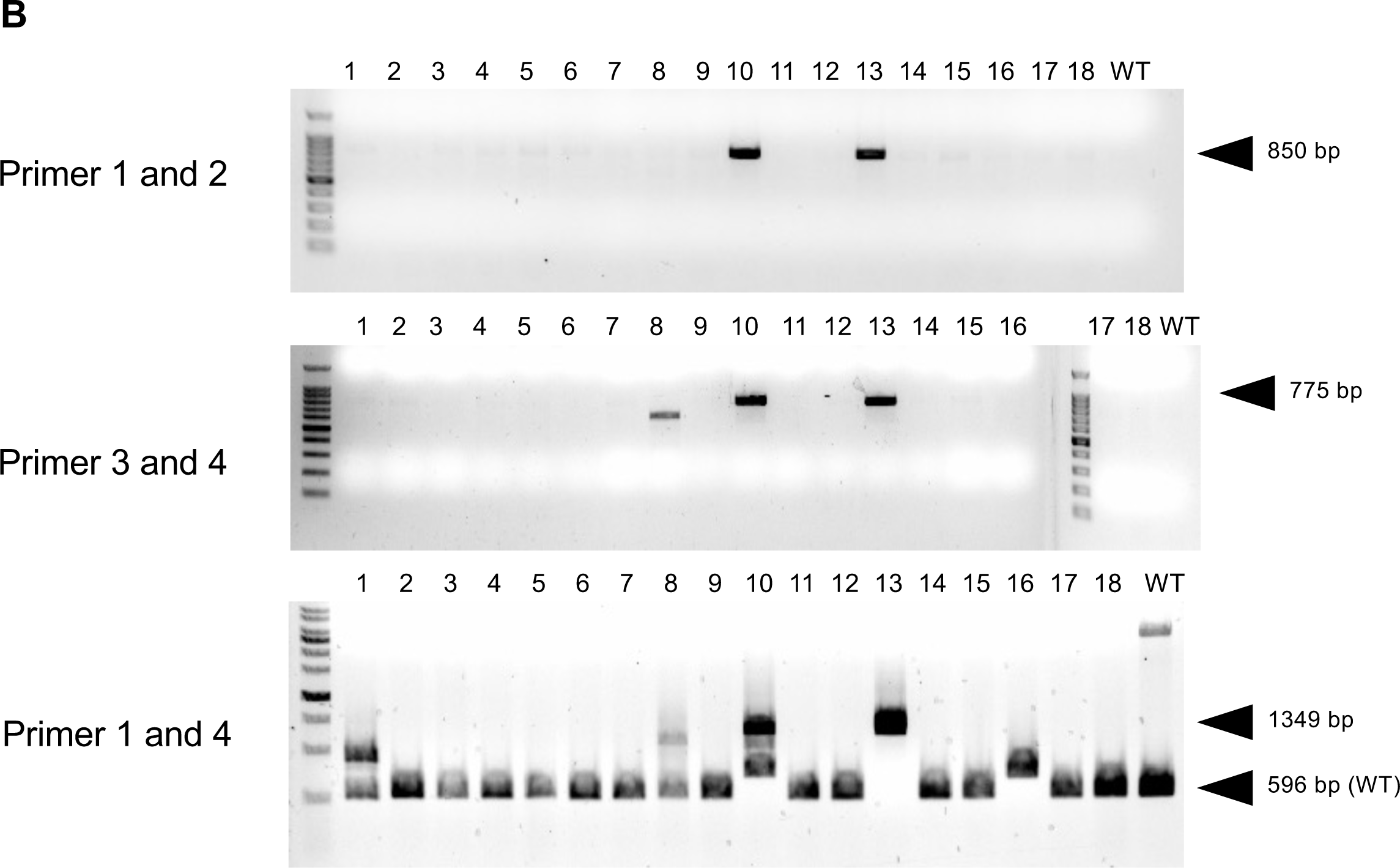
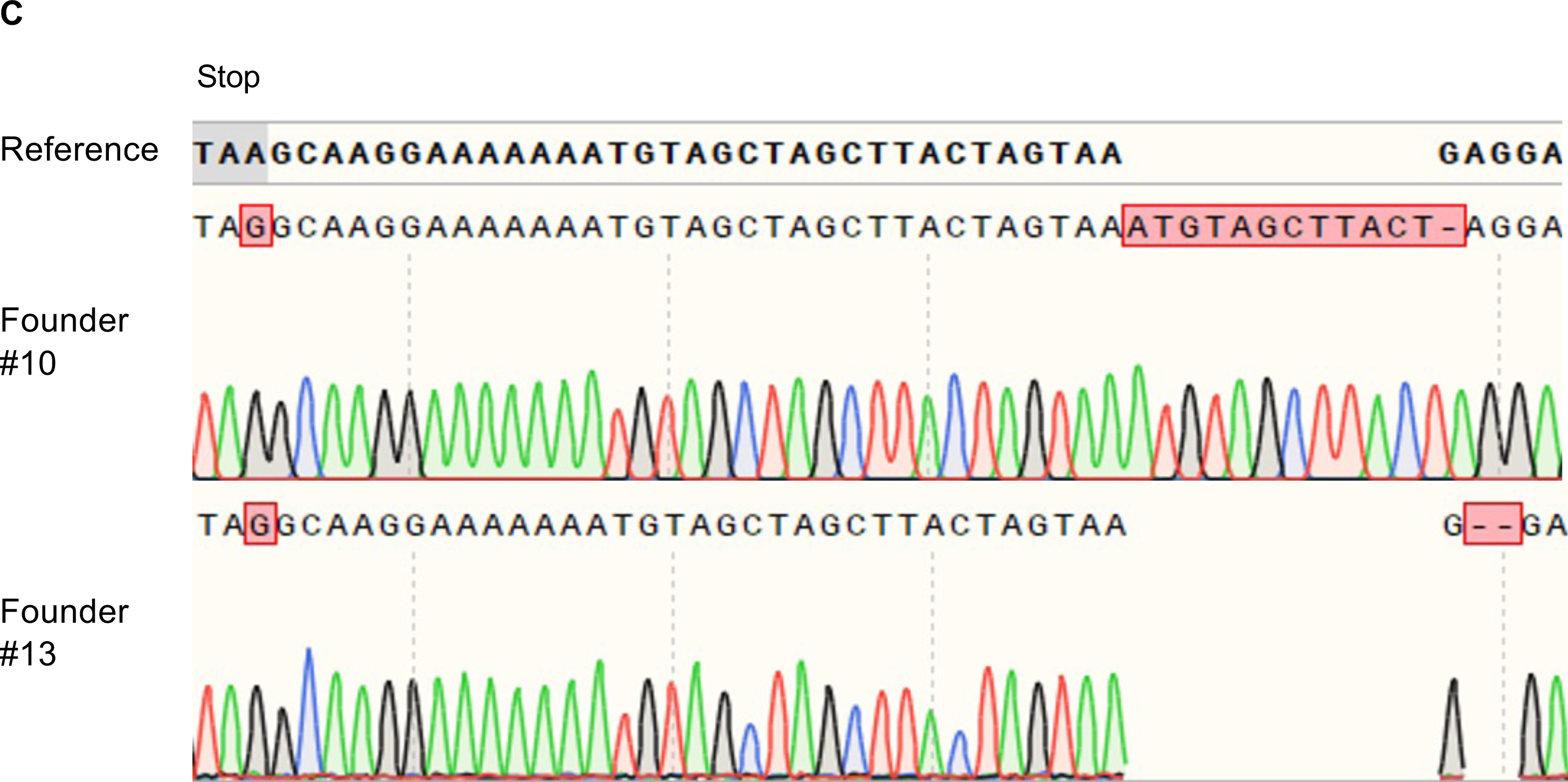
Generation of the *Rtl4*CV mice. **A. Schematic representation of the genome modification in the *Rtl4*CV mice** The Venus coding sequence was introduced into the Cas9 cleavage site upstream of the *Rtl4* stop codon together with a (GGS)x4 linker to express the RTL4-Venus fusion protein. To prevent recleavage of the *Rtl4*CV mouse genome after the genetic recombination, an A to G mutation was introduced in the codon of arginine 301 (AGG to AGA). The single-stranded donor DNA for gene modification was prepared by phosphorylated primer-mediated PCR, and the phosphorylated strand digestion with lambda exonuclease. The donor DNA has 5’ and 3’ homology arms of 161 and 221 bases, respectively. The Cas9 cleavage site was targeted near the *Rtl4* stop codon. The white squares show the 5’ and 3’ UTR in the *Rtl4* coding exon, and the gray square shows the *Rtl4* coding sequence. **B. Genomic PCR analysis of the *Rtl4*CV founder mice** The primer positions are shown in A. The founder number and wild type (WT) as a control are shown above each panel. The expected product sizes of the *Rtl4*CV allele and WT are shown to the right. **C. Genome sequences of the *Rtl4*CV founder mice** Detailed DNA sequences are shown for founder #10 and #13 with the reference genome sequence. Founder #10 and #13 have additional insertion/deletion mutations in the 3’UTR of *Rtl4*. “Stop” indicates the position of the stop codon located downstream of the Venus sequence in the *Rtl4*CV. The sequencing data were analyzed using SnapGene software.

**Fig. S2.**
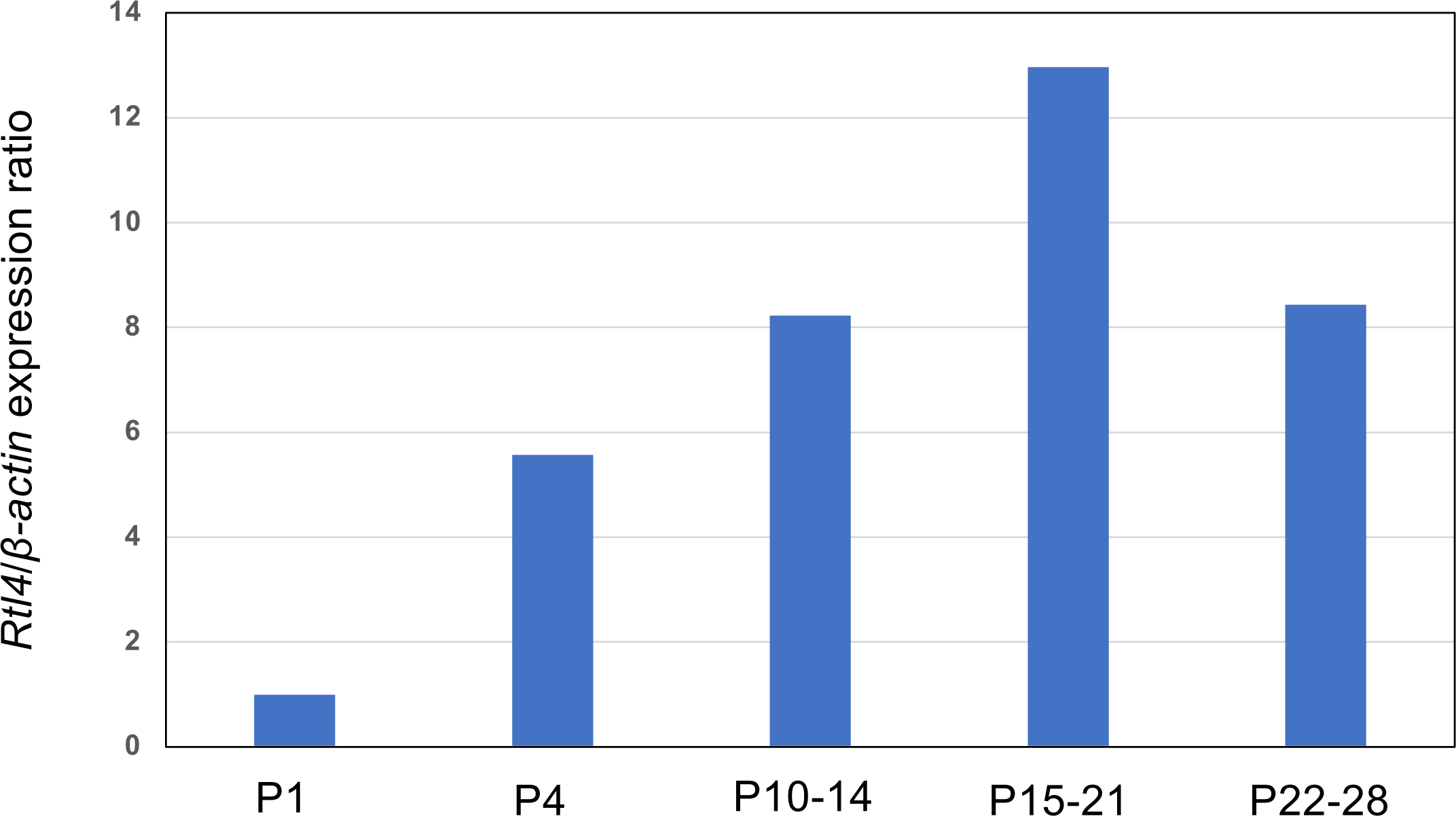
Postnatal *Rtl4* mRNA expression in WT brain. Both the *Rtl4* and *β-actin* levels were calculated with qPCR, and the ratio of *Rtl4*/*β-actin* expression was adjusted to 1 using the P1 brain level as 1. In the postnatal brain, the expression level of *Rtl4* mRNA increased gradually and peaked at 2-3 weeks. Even at P15-21, the threshold cycle (CT) of *Rtl4* was 29-30, whereas that of *β-actin* was 17-18, indicating that the level of *Rtl4* is approximately 1/4,000 of *β-actin* or less. PCR primers: *Rtl4*-F5: 5’-AAGAGGAGGATAGGAAATCACTTTG-3’ and *Rtl4*-R5: 5’-GTTGTTAGGACAAGGTTGAGG-3’; *Actb*-F: 5’-AAGTGTGACGTTGACATCCG-3’ and *Actb*-R: 5’-GATCCACATCTGCTGGAAGG-3’.

**Fig. S3.**
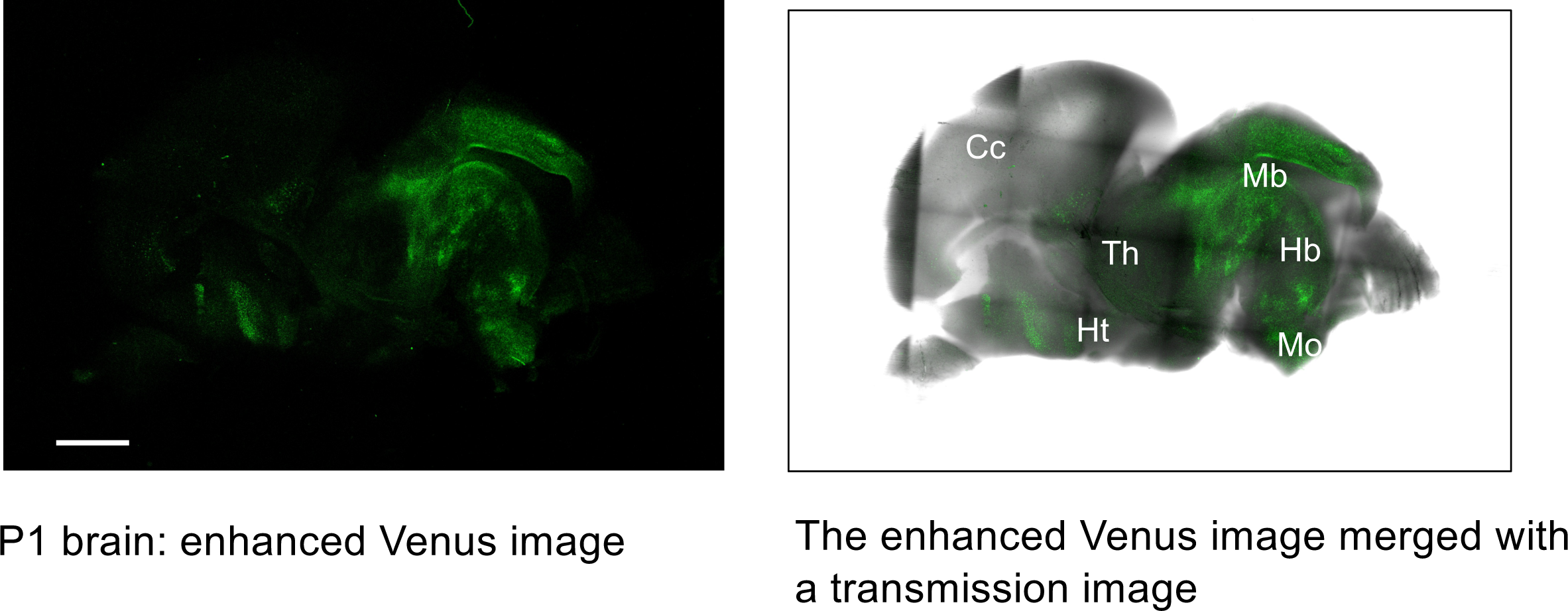
Expression of RTL4CV protein in P1 brain. RTL4CV expression was detected in certain restricted areas of the P1 brain. Left: The Venus fluorescence image. Right: The fluorescence image merged with a transmission image. Cc: cerebral cortex. Hb: hindbrain. Ht: hypothalamus. Mb: midbrain. Mo: medulla oblongata, Th: thalamus. Bar: 1 mm.

**Fig. S4.**
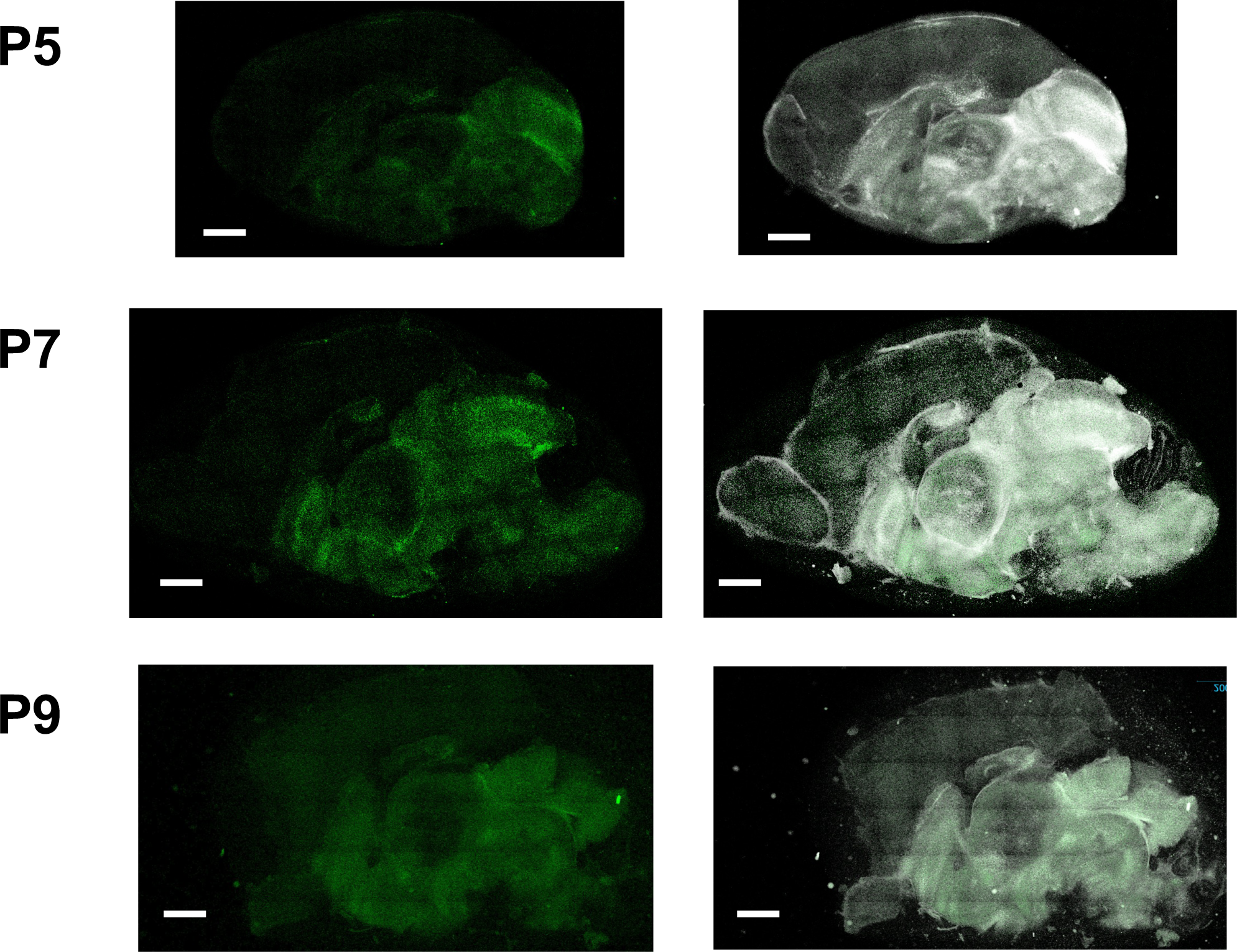
RTL4CV expression in P5-9 brain Top: In the P5 brain, a strong signal was detected in the midbrain. Middle and bottom: In the P7 and P9 brain, the hypothalamus signal became stronger, but the midbrain signal was most significant. Bar: 1 mm.

**Fig. S5.**
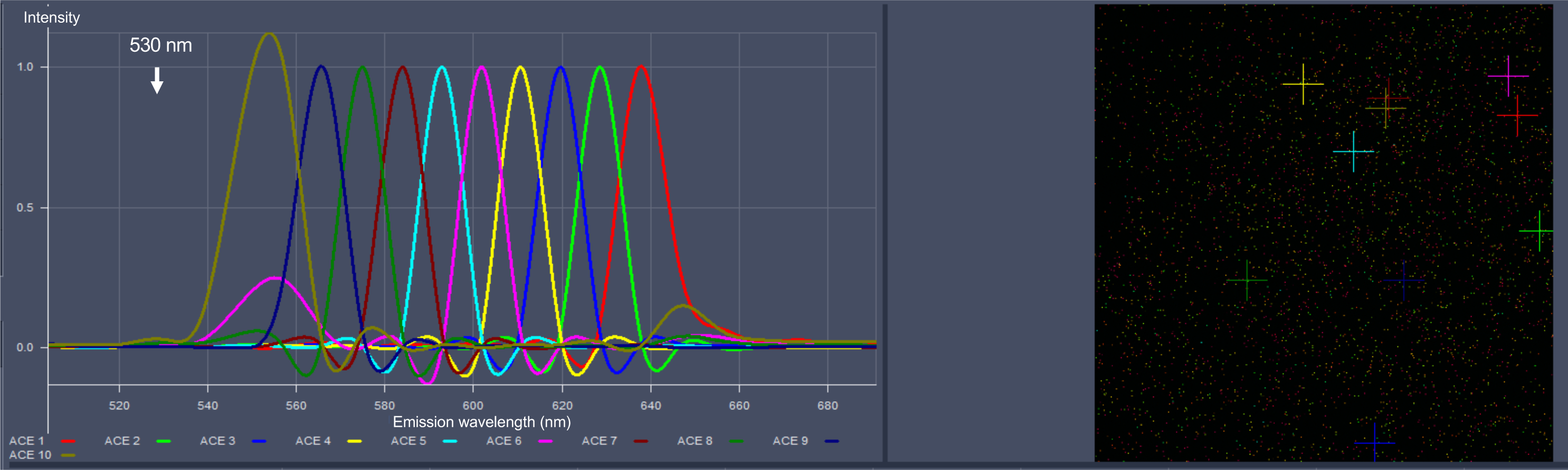
WT P21 brain analyzed by the Automatic Component Extraction (ACE) function. The hypothalamus of the WT P21 brain was analyzed using the ACE function. There is no waveform with a peak at 530 nm in the WT brain. See also the result of the *Rtl4*CV P15 brain (Fig. 1C). Left: Results from ACE1 to ACE 10. Right: The analyzed hypothalamus region. Each colored cross corresponds to the region from which the corresponding colored waveform (left) was extracted.

**Fig. S6.**
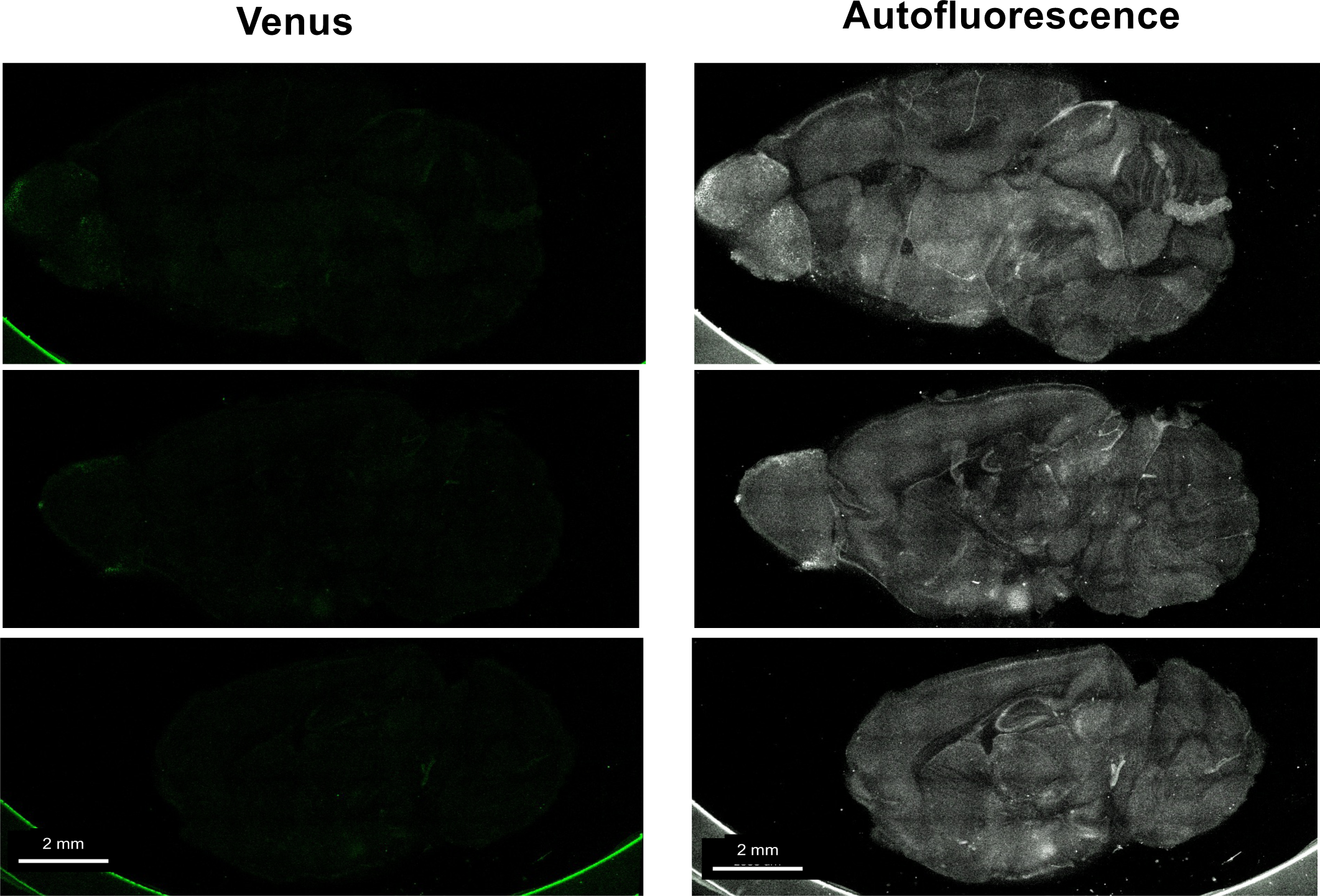
No Venus signal in WT P21 brain. In the WT brain, the waveform with a peak at 530 nm could not be detected in the top 10 (Fig. S5). The expression pattern in the WT brain was then calculated using the Venus control waveform. The signal intensity was also very low and was treated as background (BG) (see Figs. 2A and B).

**Fig. S7.**
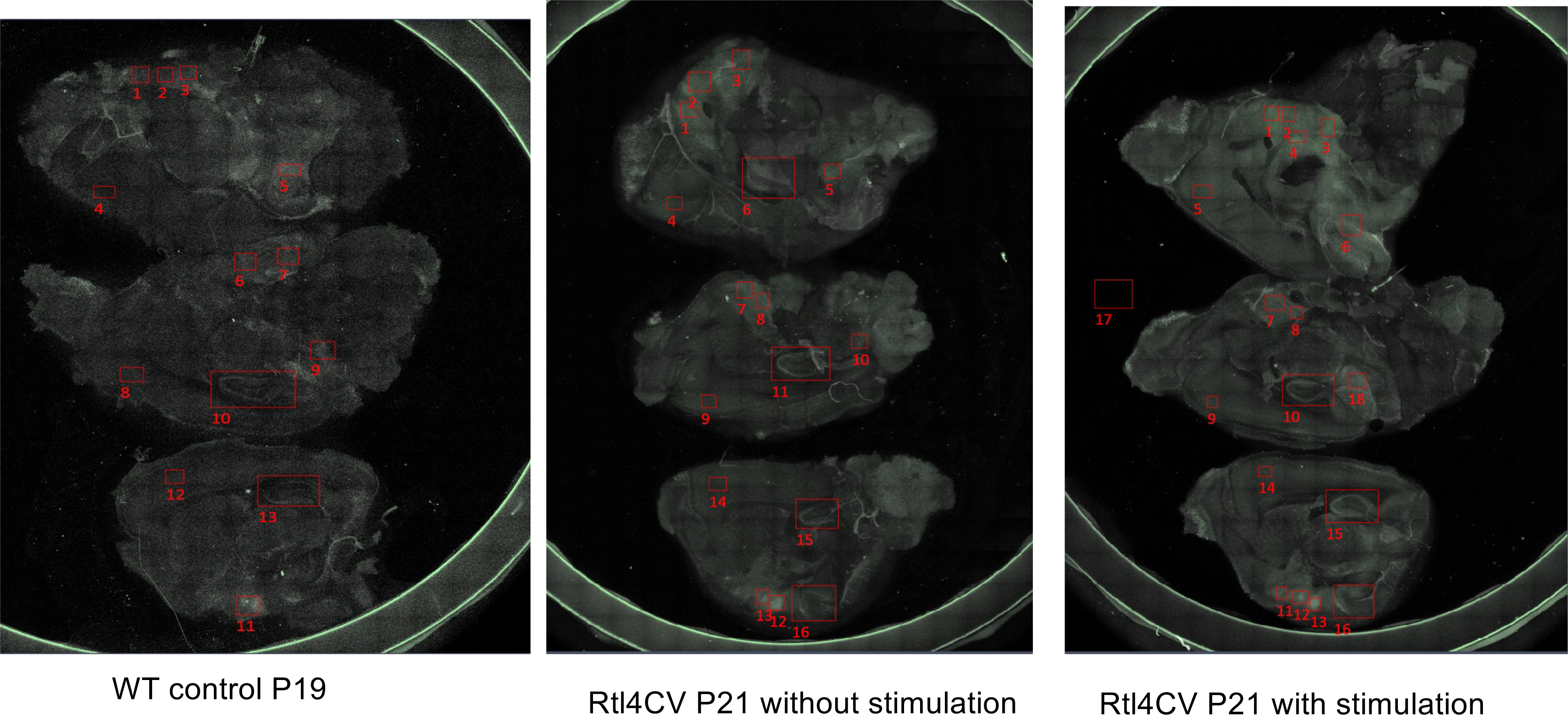
Environmental effect on the expression of RTL4CV protein in P21 brain. To calculate the Venus signal intensity in each brain region, the boxed areas were measured. The average of three to four parts in the hypothalamus and that of one to four parts in the amygdala are shown in Figure 2B. Top: Inner side of the brain hemisphere. Middle and bottom: Surfaces of brain slices of 1.5 mm width, inner (middle) and outer (bottom) images.

**Fig. S8.**
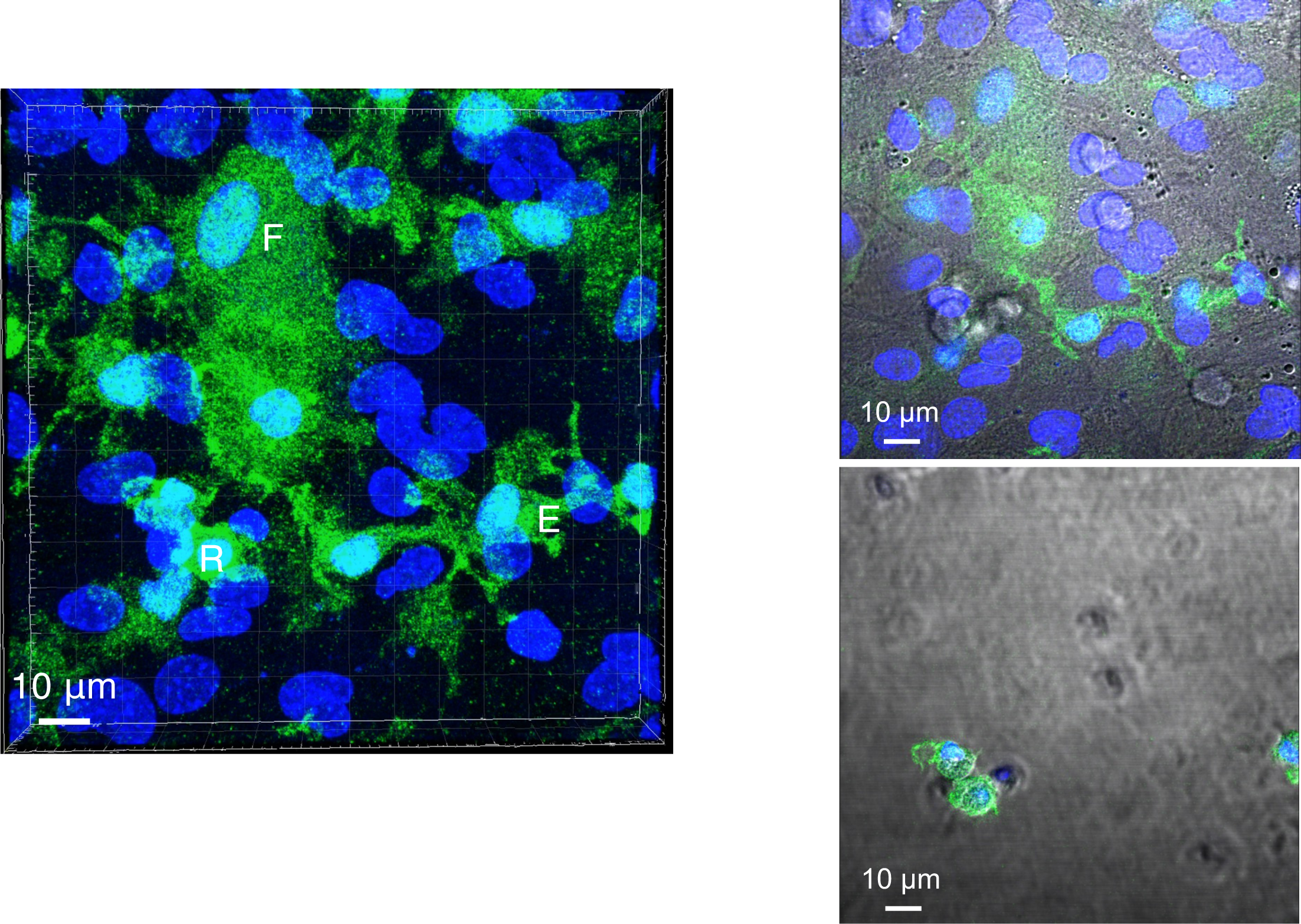
Microglial cells in the primary mixed glial culture (Iba1 staining) Several types of microglia were present in the primary mixed glial culture. Top left: flat cells below and/or within, and those with multiple processes on the astrocyte feeder layer. Bottom left: round floating cells above the feeder layer. Right: 3D movie of the primary mixed glial culture. Immunostaining with anti-Iba1-Alexa488 antibody (artificial green. DAPI (blue). Bar: 10 μm. See also 3D movie of Iba1 staining of mixed glia culture.

**Fig. S9-1.**
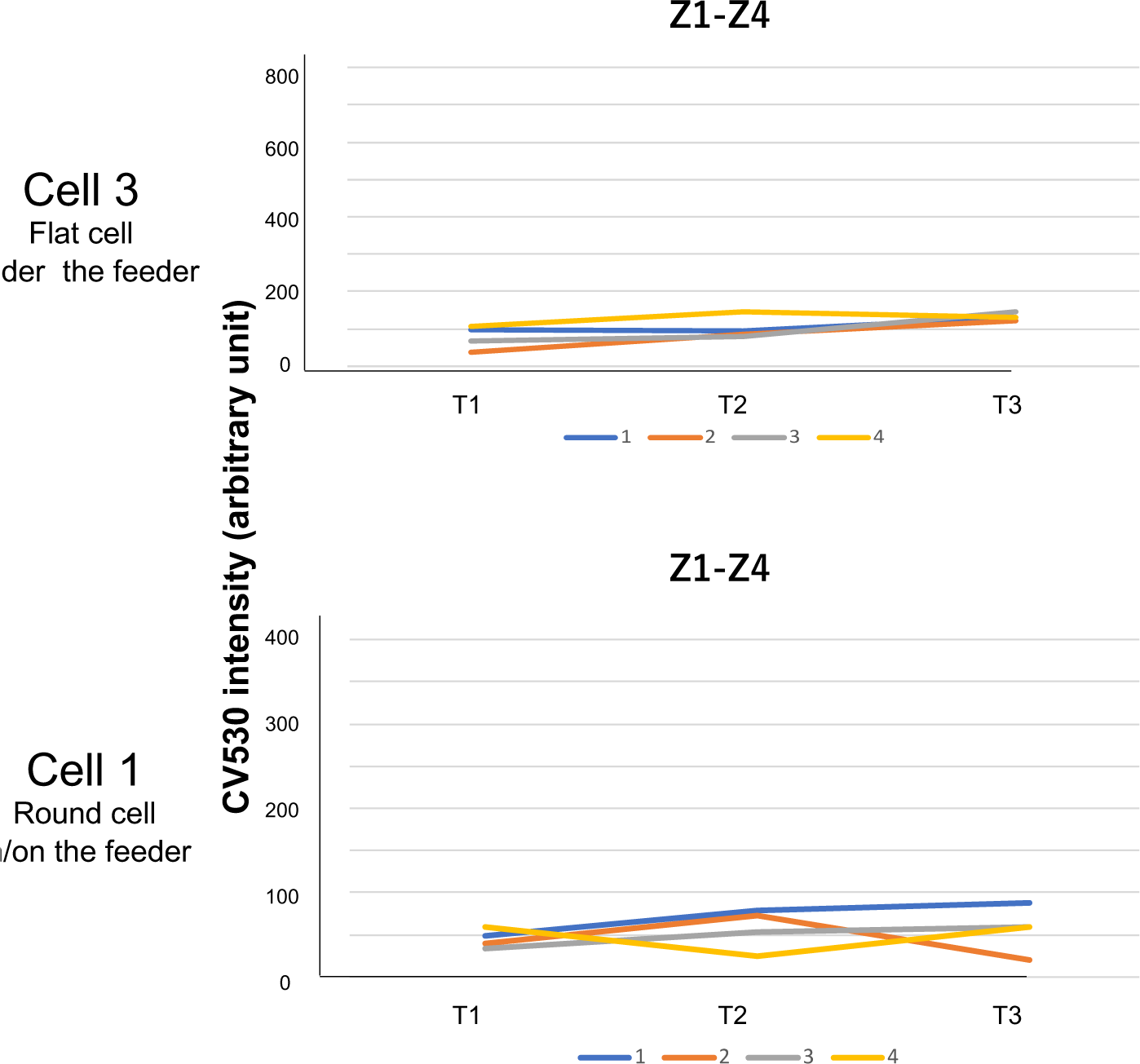
Time-lapse experiment using mixed glial culture (isoproterenol administration) The signal intensity of Z1-4 (near glass bottom) of Cell 3 (Top) and Cell 1 (Bottom) of Fig. 3C.

**Fig. S9-2.**
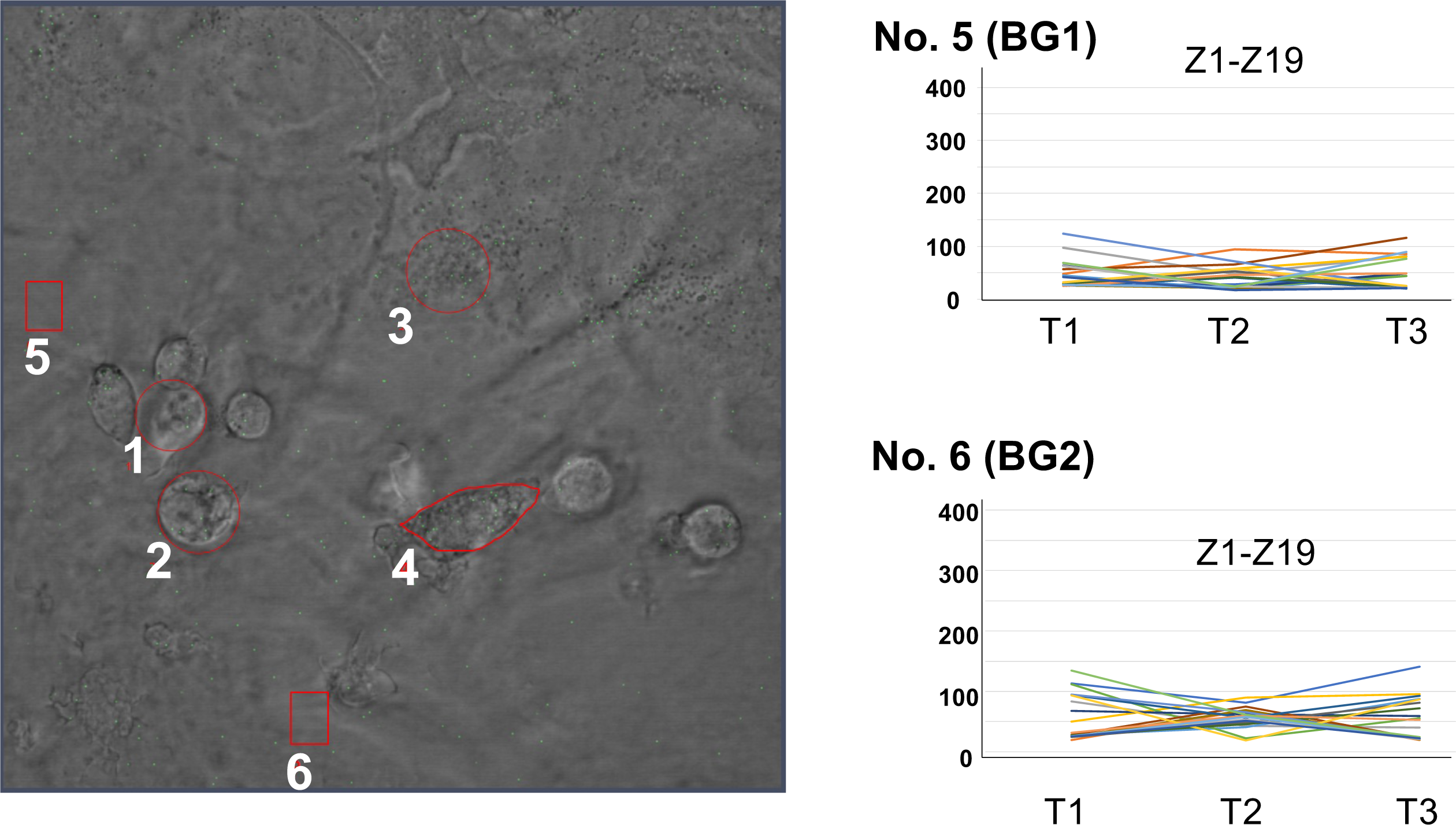
Time-lapse experiment using mixed glial culture (isoproterenol administration) Left: photo of the mixed glial culture analysed in Fig. 3C. The areas surrounded by red lines were measured: the circles (1-3) and the outlined area (4) correspond to microglial cells and the boxes (5 and 6) correspond to BG. Right: the results of the time-lapse experiment of two BG regions. The results of Cells 1 and 3 are presented in Fig. 3C and Fig. S9-1.

**Fig. S10.**
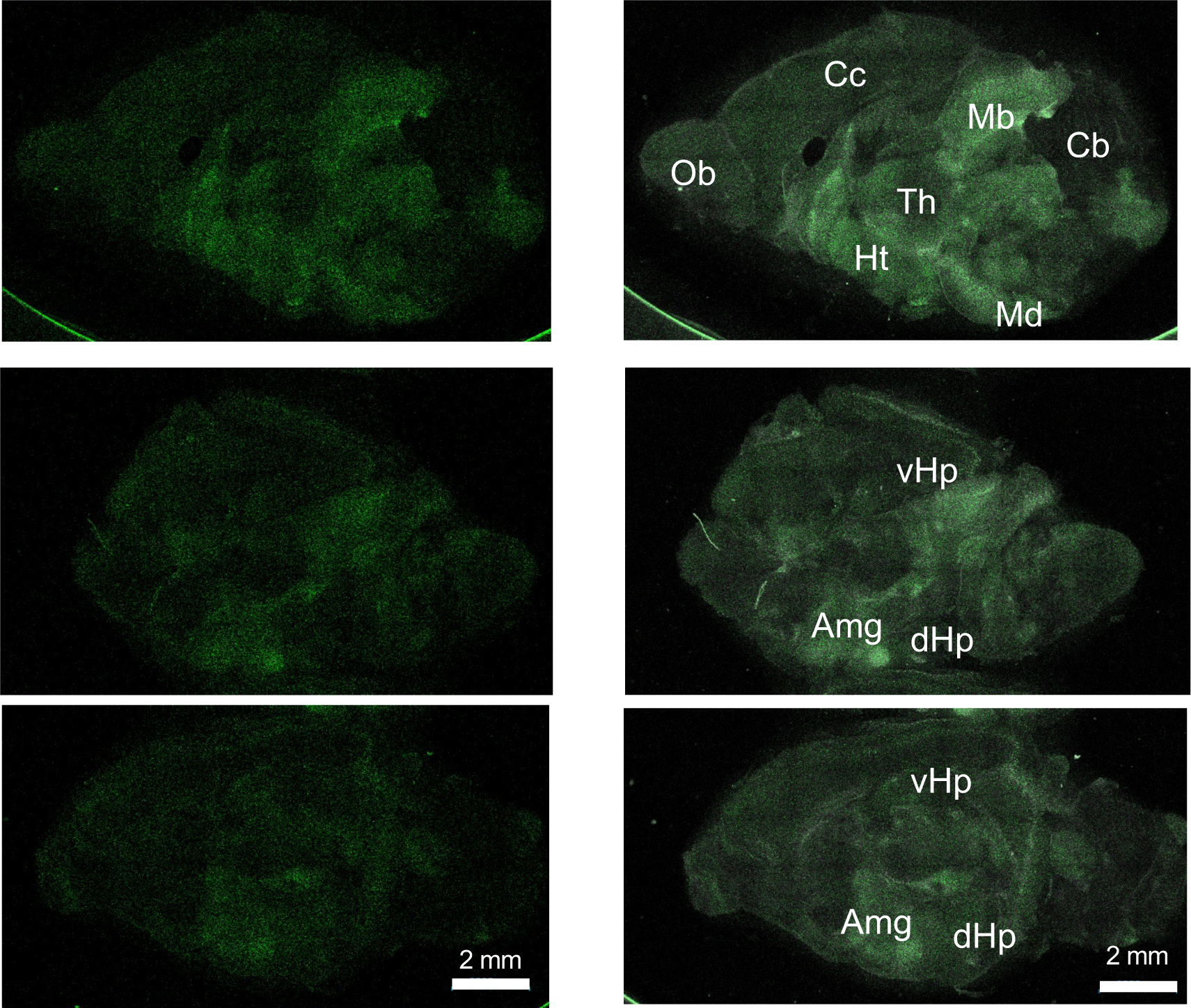
RTL4CV expression in amygdala in P10 brain. Venus images of the inner surface of the brain hemisphere (top) and brain slices of 1.5 mm width (middle and bottom). A strong Venus signal was confirmed in the amygdala as well as in the midbrain and hypothalamus (see also Fig. S4).

